# Non-linear input-output relationships in the subthalamic nucleus of Parkinson’s patients

**DOI:** 10.1101/2023.09.18.558149

**Authors:** Xiaowei Liu, Stefanie Glowinsky, Hodaya Abadi, Juan F León, Wei Wang, David Arkadir, Zvi Israel, Hagai Bergman, Jing Guang

## Abstract

Both local field potentials (LFP) and spiking (SPK) activity in the subthalamic nucleus (STN) are related to Parkinson’s disease (PD) symptoms; however, their relationship is poorly understood. We explore it by separating STN signals of 146 PD patients (308 trajectories, >25,000 recording sites) into aperiodic and periodic components and whitening these signals using their corresponding aperiodic parameters. The LFP aperiodic exponents resemble Brown noise (*α* = 2.20 ± 0.40) and are significantly higher than SPK aperiodic exponents (*α* = 0.11 ± 0.22, White noise). The periodic oscillations of LFP are overwhelmingly distributed in the high beta frequency domain while those of SPK are in both low and high beta domains. Beta oscillation center frequencies were downshifted in SPK relative to simultaneously recorded LFP. This demonstrates that the STN synaptic input (LFP) undergoes significant modifications when transformed into STN output (SPK) of PD patients, and may explain the critical role of STN in PD physiology and STN-Deep-Brain-Stimulation therapeutic efficacy.

## Introduction

Beta oscillations in local field potentials (LFP) and spiking activity (SPK) in the subthalamic nucleus (STN) are considered as the electrophysiological hallmark of Parkinson’s disease (PD)^1-7^. Bipolar LFP recordings performed within a week of electrode implantation, while patients were at rest and off medications, revealed an high proportion of patients with peaks in low beta (LBeta, 13-20Hz), high beta (HBeta, 20-35Hz) and both Beta sub-bands^4^. LBeta oscillations in LFP and spiking signals are positively correlated with the severity of PD motor symptoms, and their power is suppressed by treatment with antiparkinsonian medication or deep brain stimulation (DBS)^1, 4-6^. Moreover, studies using chronic neuronal sensing and recording devices demonstrate that beta activity is a reliable biomarker of Parkinson’s symptoms, which supports the feasibility of the personalized precision-medicine approach to adaptive neurostimulation based on the beta LFP activity^8, 9^ .

Many centers use extra-cellular recording of spiking (action-potential) activity to aid navigation to the target brain regions in DBS surgery^10, 11^. The spiking activity is a proxy for the output of the recorded neurons, and can be recorded at distances smaller than 0.1 mm from the microelectrode^12, 13^. LFP is the low frequency (e.g., 0.1-70Hz) electric potential recorded by electrodes in the extracellular space in brain tissue. LFPs are most probably generated by subthreshold (e.g., synaptic activity) modulation of the membrane potentials^14^. The exact relationship of LFPs to the neuronal activity in the STN of PD patients is still unclear. Significant coherence was found between the LFP and spiking activity in the subthalamic region^3^. Our group reported similar results in the MPTP non-human primate (NHP) model of PD^15^. Beta oscillations in LFP recordings play a role in the temporal dynamics of high frequency oscillations (HFOs) ^16, 17^. A recent study reported that periodic single-neuron bursts in the STN commonly preceded the LFP oscillation (13∼33 Hz), but that other neuronal firing activity had no relationship to the LFP^18^.

Neural oscillations have been extensively studied by advanced methods in the time and frequency domains^19-21^. The traditional oscillation bands are predefined based on the canonical frequency bands or extracted by applying narrowband filtering. Usually, the power change is implicitly assumed as a frequency-specific power change. However, most physiological phenomena follow power law (1/f, f represents the frequency) rules^22^, and the power at each frequency band is a summation of their aperiodic (1/f^α^, αis a scaling parameter that is constant over the distribution of frequencies) and periodic components. Thus, power changes of frequency bands can result from: changes in true oscillatory power, shifts in oscillatory center frequency, or changes in aperiodic parameters (offset and exponent)^23^. Extracting the periodic oscillations and aperiodic component from the signals of interest by Fitting Oscillations and One Over F (FOOOF) analysis can overcome the limitation of traditional narrowband analyses^23, 24^.

To explore the relationship between LFP and spiking activity of PD patients, we separate the STN LFP and spiking activity into periodic and aperiodic components using the FOOOF algorithm^23^, and whiten the neuronal activity using the aperiodic exponent.

## Results

Electrophysiological recording of the STN activity and neighboring structures was done as part of the standard-of-care DBS navigation procedures. All signals were recorded when the patients were awake and in a state of rest. The LFP and spiking activity were obtained by offline filtering the raw data at 3-200Hz and 300-6000Hz respectively, using 4 poles Butterworth, zero-phase band-pass filters. The spiking activity was rectified^1^ to reveal the low-frequency (<300Hz) oscillations in discharge rate (Fig. 1 and S1). Based on our inclusion criteria, we included 308 out of 492 trajectories from 146 patients, and 25,822 and 27,130 recording sites of LFP and spiking activity, respectively. Further details are shown in Table 1. The FOOOF algorithm^23^ decomposed the neuronal activity into aperiodic and periodic components (Fig. 1). The aperiodic exponents were used to whiten the power spectral densities (PSDs) of the LFP and the rectified spiking discharge rate (SPK) activity.

**Table 1.** Demographics of patients, trajectories and STN.

| **Table 1. Demographics of patients, trajectories and STN** |  |  |
| --- | --- | --- |
| **Demographics** | **Results** |  |
| Patients (N) | 146 |  |
| Age (years) (Mean ± SD) | 62.03 ± 9.63 | |
| Gender (N, %) |  |  |
| Male | 100 (68.49%) |  |
| Female | 46 (31.51%) |  |
| Disease Duration (years) (Mean ± SD) | 10.17 ± 3.84 | |
| Preoperative LEDD (mg) (Mean ± SD) | 1,020.71 ± 545.48 | |
| Preoperative UPDRS-□ scores (Mean ± SD) | | |
| Off medication | 43.43 ± 12.73 | |
| On medication | 19.76 ± 8.99 | |
| Total trajectories (N) | 492 |  |
| Trajectories included (N) | 308 |  |
| Right (E1, E2) | 156 (68, 88) |  |
| Left (E1, E2) | 152 (72, 80) |  |
| Recording Sites | LFP | SPK |
| Total sites (N) | 42,680 | 42,680 |
| Sites included (N, %) | 25,822 (60.50%) | 27,130 (63.57%) |
| Sites excluded (N, %) | 16,858 (39.50%) | 15,550 (36.43%) |
| Short signal length (N, %) | 1,288 (3.02%) | 1,288 (3.02%) |
| Outliers of RMS (N, %) | 2,262 (5.30%) | 434 (1.02%) |
| Trajectories excluded (N, %) | 13,308 (31.18%) | 13,828 (32.40%) |
| The length of sub-regions (mm) (Mean ± SD) | LFP | SPK |
| Pre-STN | 4.35 ± 1.51 | 4.40 ± 1.50 |
| DLOR | 3.03 ± 1.08 | 3.06 ± 1.09 |
| VMNR | 2.63 ± 0.96 | 2.63 ± 0.96 |
| DLOR-Per | 0.53 ± 0.14 | 0.53 ± 0.14 |
| STN: Subthalamic nucleus. SD: Standard deviation. N: number. LEDD: levodopa equivalent daily dose. E1: electrode 1 (Ben-Gun posterior location). E2: electrode 2 (Ben-Gun central location). DLOR: Dorsal lateral oscillatory region. VMNR: Ventral lateral non-oscillatory region. DLOR-Per: The percentage of DLOR length out of the total STN length (DLOR+VMNR) |  |  |

**Figure 1:**
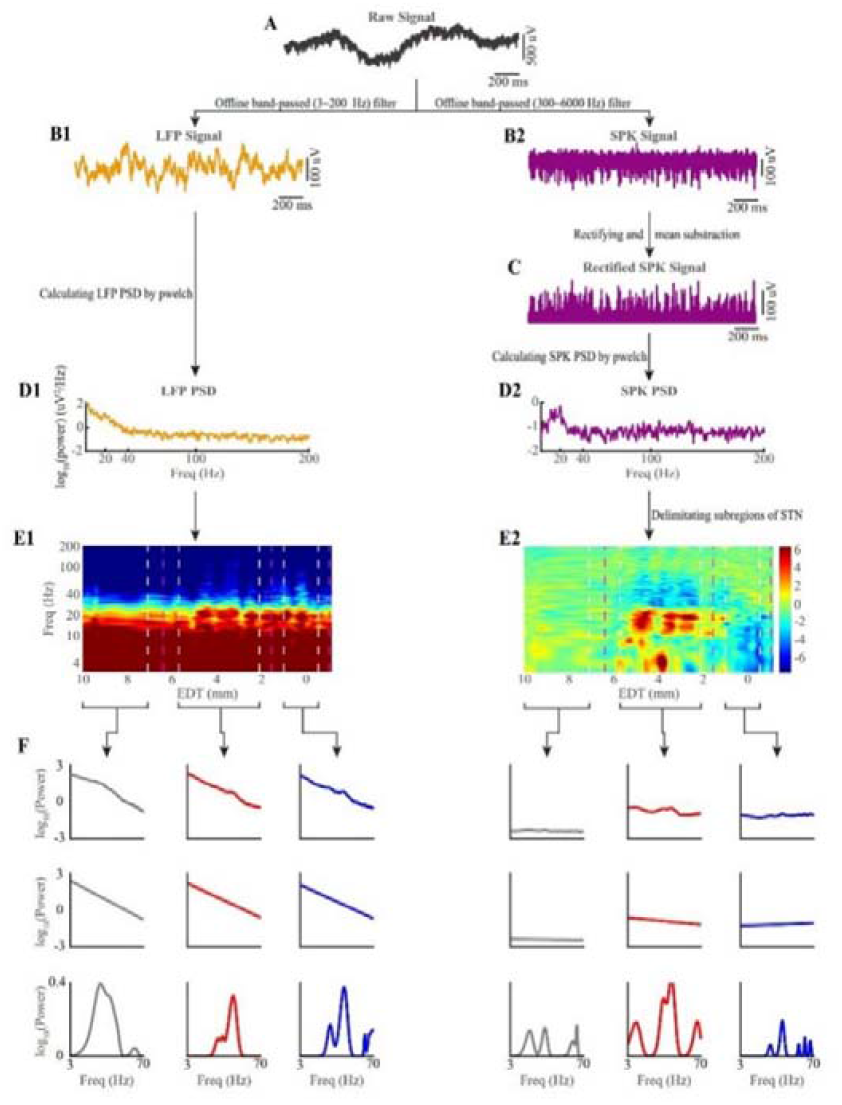
Electrophysiological pre-processing and analysis. (A) Two seconds’ example of raw signal band-passed filtered from 0.07 to 9000 Hz. (B1) The LFP signal is obtained by zero-phase band-pass filtering the raw signal from 3-200Hz. (B2) The spiking (SPK) signal is obtained by zero-phase band-pass filtering of the raw signal from 300 to 6000 Hz. (C) The rectified SPK signal is obtained by applying the absolute operator to the spiking signal and then subtracting the mean. (D1) The LFP power spectral density (PSD) in the range of 3 to 200 Hz is obtained by pwelch function (MATLAB). (D2) The SPK PSD (3 to 200 Hz) is obtained by applying the pwelch function to the mean-subtracted rectified SPK signal. (E1 and E2) Spectrograms and delimitating sub-regions of STN. X-axis is the estimated distance to the target (EDT). Y-axis is the frequency from 3 to 200 Hz (logarithmic scale). The spectogram’s color-scale represents 10*log_10_(spectral power / average spectral power). The first and third vertical magenta dashed lines indicate the entry and exit of STN, respectively, and the second one represents the boundary between the dorsolateral oscillatory region (DLOR) and the ventromedial non-oscillatory region (VMNR). The vertical white dashed lines represent the safe 0.5 mm margins of each sub-region. STN borders were found by hidden Markov analysis (HMM) of neural spiking in E2, and were copied to the same trajectory LFP data in E1. (F) The first row shows the averaged PSD (from 3 to 70 Hz) of each sub-region of a single trajectory. The aperiodic (row 2) and periodic (row 3) components of LFP and SPK activity of this single trajectory are obtained by applying FOOOF analysis to the averaged PSD in the first row. The columns indicate the regions: pre-STN (grey), STNDLOR (red) and STN VMNR (blue). See also Figure S1.

### Goodness of fit of the FOOOF analysis to the LFP and SPK activity

We applied the FOOOF algorithm to both LFP and SPK single site recordings to separate their PSDs into aperiodic and periodic components. Fig. 2 depicts the STN LFP and SPK population mean of the raw PSDs and their aperiodic and periodic components. The goodness of fit of the FOOOF analysis is assessed by the R^2^ and mean absolute error values (MAE, error). Optimally, R^2^ and error should be as close as possible to 1 and zero, respectively. The R^2^ values of LFP are 0.99 ± 0.01 (mean ± SD) in the three STN subregions. The R^2^ values of SPK are 0.64 ± 0.17, 0.89 ± 0.15 and 0.65 ± 0.17 in Pre-STN, dorsal lateral oscillatory region of STN (DLOR) and ventral medial non-oscillatory region of STN (VMNR), respectively.

**Figure 2:**
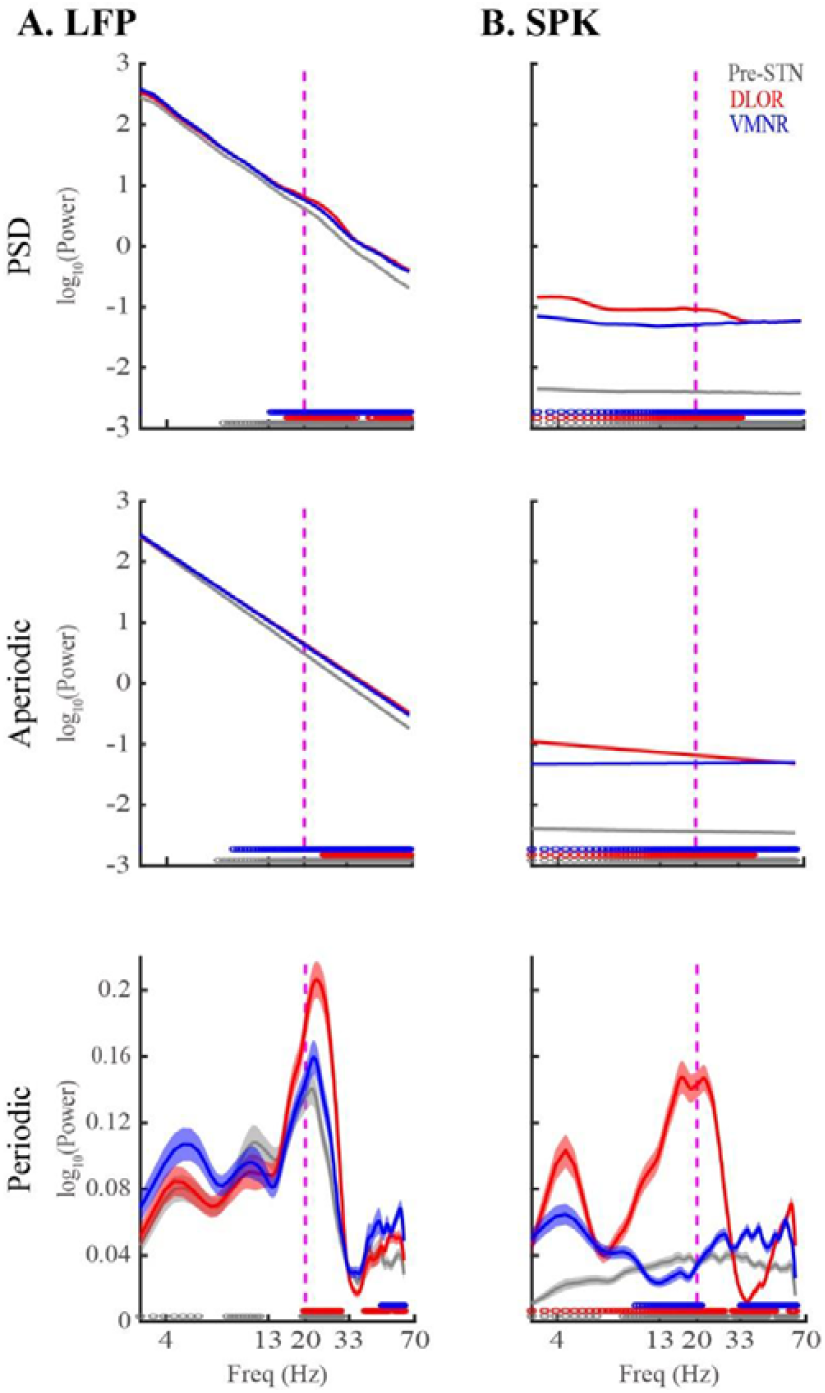
Robust differences in aperiodic and periodic components of subthalamic LFP and spiking (SPK) population activity. (A) Population mean LFP PSD (row 1) and its aperiodic (row 2) and periodic (row 3) components in three STN sub-regions. (B) Population mean SPK PSD (row 1) and its aperiodic (row 2) and periodic (row 3) components in the same three STN sub-regions. The grey/red/blue lines indicate the pre-STN, STN-DLOR and STN-VMNR, respectively. Their corresponding shade lines indicate SEM. The colored circles above the x-axes represent the frequencies at which there was a significant difference between the pre-STN and DLOR (grey), between the DLOR and VMNR (red), and between the VMNR and pre-STN (blue). Significance was calculated using the Wilcoxon rank sum test and the Bonferroni correction (p < 0.05/3 = 0.0167). Vertical dashed lines represent the 20 Hz frequency point. See also Figures S2, S3, S4, S5 and S6.

The SPK lower R^2^ values can be explained by its lower exponent values relative to LFP (Fig. 2 and 3B). Our numerical simulations (Figure S2A and S2B) demonstrate that exponents close to zero (as for our SPK activity) yield lower R^2^ values, and that as the absolute value of the exponent grows, the R^2^ value approaches 1. The addition of the periodic component of the signal reduces the effect of the exponent on the R^2^ values. This explains why the R^2^ values of SPK in pre-STN and VMNR are lower than those in the DLOR with more prominent periodic components (Fig. 2 and S2). Notably, the error is not affected by the span of the periodic power and exponent (α) values (Fig. S2C). Indeed, both LFP and SPK have low error values in three STN subregions (error < 0.04). We concluded that the FOOOF analysis provided a good fit for our LFP and SPK data in the three STN subregions, and proceeded to compare their aperiodic and periodic components.

**Figure 3:**
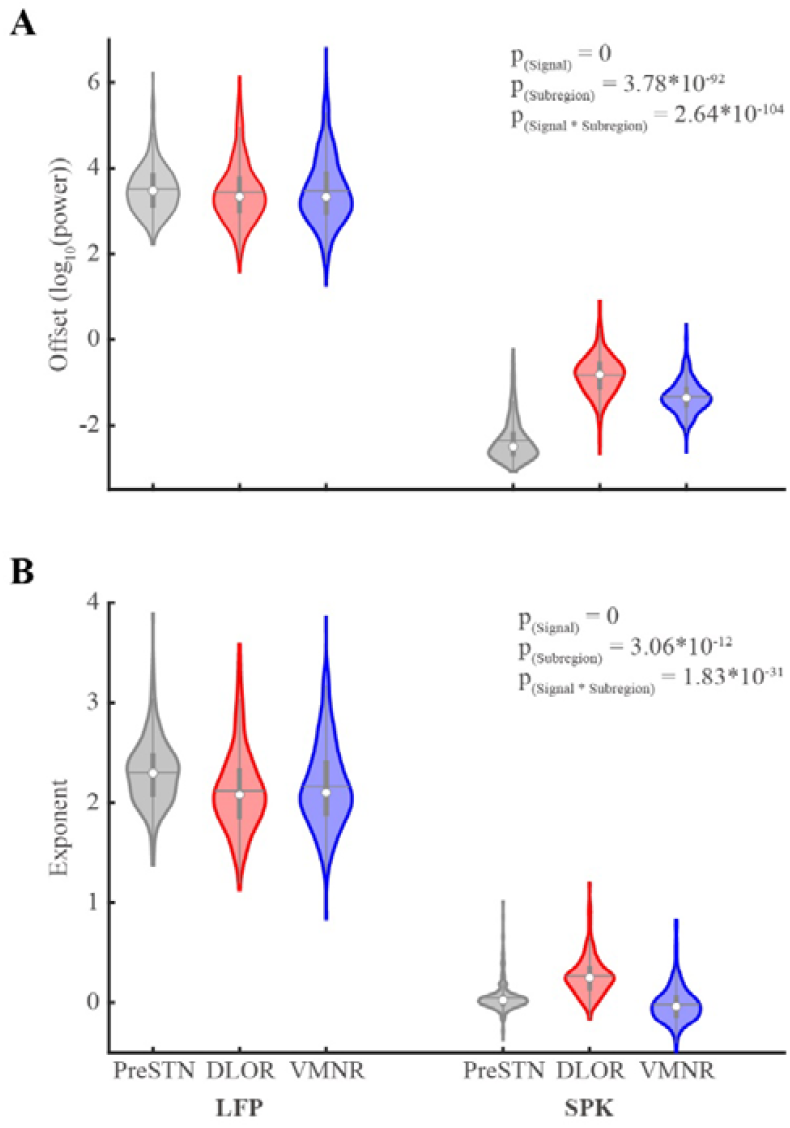
Significant differences in aperiodic parameters of LFP and spiking (SPK) activity in the sub-regions of the subthalamic nucleus. (A) The aperiodic offset parameter of LFP and SPK in three STN sub-regions. (B) The aperiodic exponent parameter of LFP and SPK in three STN sub-regions. The pre-STN is shown in grey, the STN-DLOR in red, and the STN-VMNR in blue. The contour of the violin plots shows the distribution of the data. The white circle shows the median. The horizontal grey line represents the mean. The grey vertical bold lines span from the 25^th^ to the 75^th^ percentiles of the sample, and the length of this line is the inter-quartile range. The lowest and highest whiskers of the violin plots are values which are 1.5 times the inter-quartile range below the 25^th^ percentile and above the 75^th^ percentile. The N-way analysis of variance was used to analyze the difference of aperiodic parameters. The P_(Signal)_ indicates that there is a significant difference in offset or exponent values between LFP and spiking (SPK) activity (offset: p = 0; exponent: p = 0). The P_(Subregion)_ indicates the statistical difference of offset or exponent values between pre-STN, DLOR and VMNR (offset: p = 3.78*10^-92^; exponent: p = 3.06*10^-12^). The P_(Signal*Subregion)_ represents the interaction effect of the difference of offset and exponent values between the signal types and sub-regions (offset: p = 2.64*10^-104^; exponent: p = 1.83*10^-31^). Detailed results of multiple comparisons are shown in Table S1. See also Figures S2, S3, S4, S5 and S6.

### Significant differences in aperiodic parameters between LFP and SPK activity

The aperiodic parameters are offset and exponent^23^. LFP has much larger offsets than SPK in the three subregions (Fig. 2 and 3A). There is no significant difference in LFP offsets between subregions, while SPK offsets in each subregion are similar but significantly different (Table S1).

The exponents of LFP and SPK are significantly different (2.20 ± 0.40 and 0.11 ± 0.22, respectively, Fig. 3B). The exponents of LFP and SPK resemble those of Brown noise (α=2) and White noise (α=0), respectively (Fig. 2 and 3B). LFP exponent in the pre-STN is significantly larger than that in the two subthalamic regions. The detailed results of our multi-comparison analysis are shown in Table S1.

### Positive correlation between aperiodic parameters of LFP and SPK activity

We evaluated the relationship between aperiodic parameters of LFP and SPK. We found a robust and significant correlation between aperiodic parameters within a signal type (i.e., LFP or SPK) in Pre-STN, DLOR and VMNR. The positive correlation between the offset and exponent in both LFP and SPK signals is strongest in DLOR (Fig. S3). Additionally, we found a mild correlation between different signals’ aperiodic parameters in DLOR (Fig. S3). This suggests that the relationship between these parameters is specific to each signal type and does not generalize across different signals in Pre-STN and VMNR, which is different from that in DLOR.

### Robust differences in periodic power between LFP and SPK activity

Fig. 2 bottom subplots depict the population periodic activity of LFP and SPK in Pre-STN, DLOR and VMNR. Beta oscillations in LFP are clearly observed in the three subregions (Fig. 2A), while in SPK they only exist in DLOR (Fig. 2B). LFP HBeta (20-33Hz) oscillation in DLOR is significantly higher than that in both Pre-STN and VMNR. There is no significant difference in LFP LBeta oscillations between the three subregions.

SPK has smaller beta power than LFP. Additionally, the frequency distribution of beta oscillations in SPK is obviously different from that in LFP (it is shifted left relative to the LFP). LFP also shows theta and alpha oscillations in the three subregions (Fig. 2A), but SPK demonstrates theta oscillations in DLOR and no robust oscillations in Pre-STN and VMNR (Fig. 2B).

### The difference between LFP and SPK periodic and aperiodic results is not due to the data processing methods

To verify that the differences of aperiodic and periodic components between LFP and SPK are not an artifact of the rectification of the SPK signal (Fig. 1C), we also applied rectification to the LFP (Fig. S4). The rectified LFP has a strong goodness of fit of FOOOF analysis (R^2^∼0.99, error<0.05; Fig. S4C and S4D). The aperiodic parameters of non-rectified and rectified LFPs don’t reveal significant qualitative differences (Fig. S4C and S4D). The rectified LFP offsets (Table S2 and Fig. S4C and S4D) are statistically non-different between the three subregions and are much larger than those of SPK (Fig. 3A). The rectified LFP exponent in Pre-STN is steepest, which is same as the non-rectified LFP (Table S3, Fig. S4C and S4D). The average exponent of rectified LFP in the three subregions is 1.78 ± 0.42, which still resembles Brown noise.

There are no significant qualitative differences between non-rectified and rectified LFP in both their PSDs (frequency range>13Hz) and aperiodic power (Fig. S4A and S4B). Rectified LFP has higher periodic power in theta and alpha frequency bands, compared with the non-rectified LFP (Fig. S4A and S4B). This is in line with previous studies demonstrating that full-wave rectification of EMG demodulates and enhances underlying low-frequency components of the signal (“carrying” frequencies), which may not be observed in the original signal due to the greater power of higher-frequency components of the signal^25^.

We used a numerical simulation to further verify that our observed shift in center frequency of beta oscillations in SPK is not an artifact of our data processing. We simulated Brown noise signals to which we added beta modulation and spikes (Fig S5). The simulation demonstrates that LFP rectification smooths the power distribution in the beta region, but doesn’t change the center frequency (Fig. S5A-D). After the addition of Poisson distributed spikes following a threshold crossing, the offset and the exponent of the simulated LFP don’t change (Fig. S5F and S5G). After the band-pass (300-2000 Hz) filtering of the wide-band signal in figure S5G, the remaining signal lost its low-frequency components (Fig. S5H). However, the full-wave rectification reinstated the low-frequency (20 Hz) oscillatory component (Fig. S5I). These results reveal, in line with our previous studies^26^, that spikes don’t affect the LFP behavior, and that rectification (absolute operator) of the spiking (>300Hz) activity expose the behavior of the discharge rate of the spikes.

Fig. S6 shows the differences between the envelope of the discharge rate (SPK, as used in the DBS physiological navigation algorithms^27, 28^ and in this study) versus the analog broad band (3-9000Hz) neuronal activity that include both the LFP and the extracellularly recorded raw spiking activity. The broad-band neuronal signal can be represented by power law distribution with exponent values higher than 2. However, even though having good fitness (Fig. S6B), such broad-band presentation of the neural activity masks the low-frequency oscillations that characterize the LFP and the discharge rate of the STN in the parkinsonian state. This demonstrates the importance of analyzing the LFP and SPK signals separately, as is done in the remainder of this paper.

### Different distribution of beta oscillation in DLOR of STN between LFP and SPK raw and whitened spectrograms

To reveal the periodic activity, we used the FOOOF exponents to whiten the signals (in frequency and time domain) at the level of each recording site (Fig. S1). The average population spectrograms in fig. 4 are whitened in frequency domain. The whitened LFP average spectrogram demonstrates more clear oscillations than the raw LFP (Fig. 4A, LFP). Robust LFP HBeta periodic activity in the STN becomes visible in the whitened spectrogram after the removal of activity resulting from volume conductance (e.g., from cortical activity). We did this using z-score normalization based on the pre-STN activity for each frequency bin (Fig. 4B, LFP).

**Figure 4:**
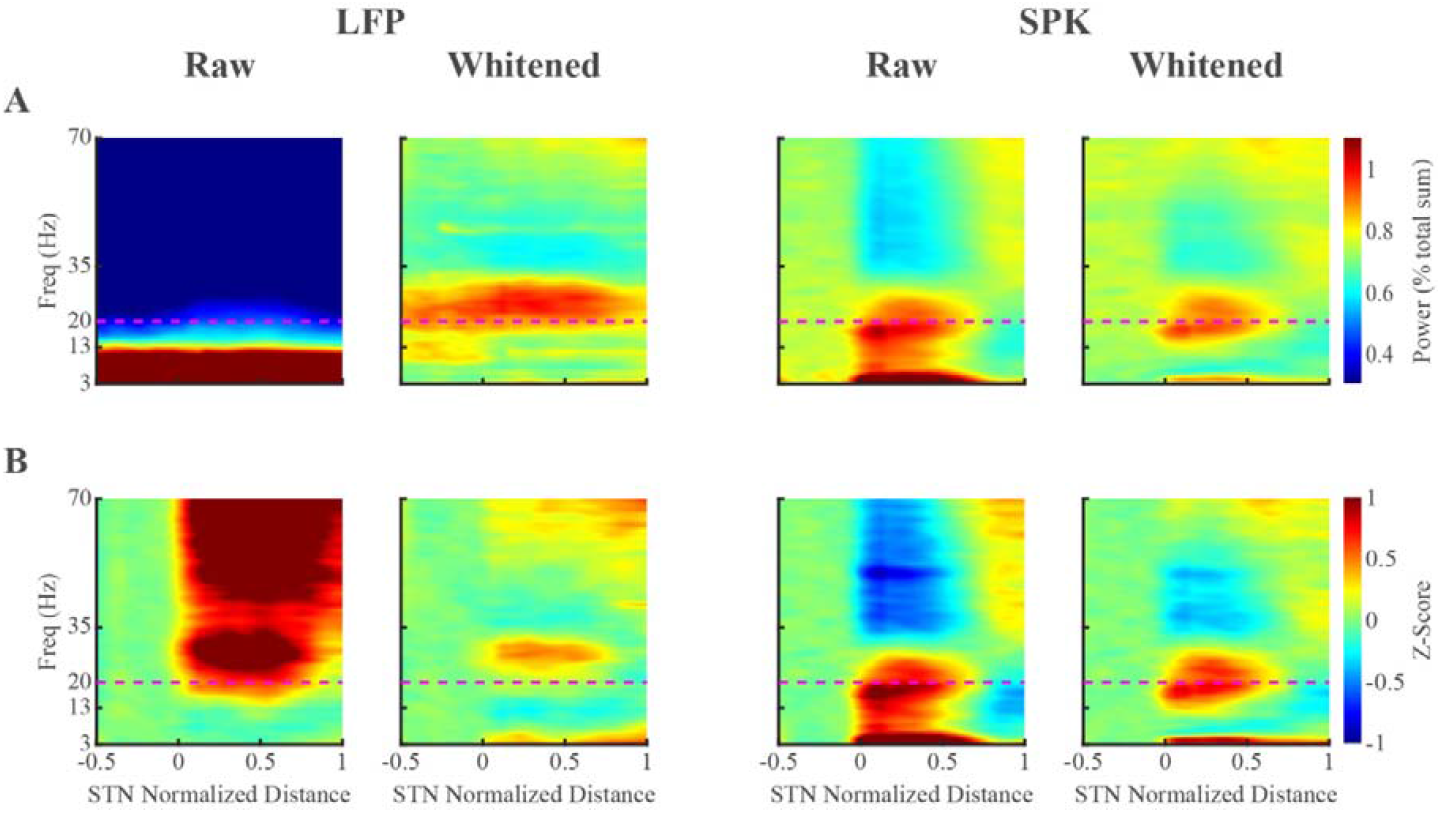
Raw and whitened averaged spectrograms reveal differences in beta frequency distribution between LFP and spiking (SPK) activity in the dorsolateral oscillatory region of subthalamic nucleus. (A) Raw and whitened spectrograms of LFP and SPK are normalized by the total amount of power in the tested frequency range (3-70Hz) for each tested distance site (normalization by frequency). (B) The raw and whitened spectrograms of LFP and SPK are normalized by frequency (as in A) and by the power in the pre-STN domain per each frequency bin (normalization by distance). The spectrograms in columns 2 and 4 of A and B are whitened in frequency domain (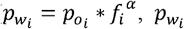 is the whitened power at the i^th -^ frequency, 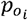 is the original power at the i^th^ frequency, *f*_*i*_ is the i^th^ frequency, and α is the aperiodic exponent of each recording site). The x-axis is the normalized distance to the target (normalize STN length from entry to exit to 1). The entrance and exit of STN are represented by 0 and 1, respectively. The negative values on the x axis indicate the pre-STN region. The y-axis is frequency in linear scale. The color-scale of the power spectral density normalized by frequency (A) indicates the percentage power of frequency bin out of total power. The color-scale of the power spectral density normalized by frequency and by distance (B) represents the deviation from the mean value of the first 10 depths in pre-STN (z-score, standard deviation unit). The horizontal magenta dashed line is the referenced line of 20 Hz. See also Figure S7.

In sharp contrast with the LFP, the whitened SPK spectrogram displays both LBeta and HBeta oscillations in DLOR of STN (Fig. 4A and 4B). Thus, we found that LFP and SPK have different distributions of beta oscillations in DLOR of STN. To verify that this isn’t due to the frequency domain whitening procedure, we compared the results of the classical frequency domain method and the time domain whitening method, and found them to be similar in both cases (Fig. S7).

### Lower peak beta frequency in SPK relative to LFP in the average population PSDs of DLOR of STN

Following the analysis of the different frequencies presenting in whitened SPK versus whitened LFP spectrograms (Fig. 4), we analyzed their whitened PSDs in Pre-STN, DLOR and VMNR separately (Fig. 5). In DLOR of STN, both whitened LFP and whitened SPK have clear beta oscillations, though whitened LFP has peak beta power in HBeta, while whitened SPK is in the LBeta range (Fig. 5A). In both Pre-STN and VMNR, the alpha and beta oscillations appear in whitened LFP, but not in whitened SPK. We applied z-score normalization based on the pre-STN activity to reduce confounding effects of volume conductance on the STN activity. The locations of the peak beta power in whitened LFP and whitened SPK in DLOR are still above and below the 20 Hz HBeta-LBeta division line, respectively (Fig. 5B). Whitened LFP has a higher frequency of peak beta power after this normalization compared to before (Fig. 5). The beta oscillations in whitened LFP in VMNR of STN may partly come from the DLOR, because the detection of the transition between the DLOR and VMNR is less accurate and the boundary between the two subthalamic subregions is not always sharp (sometimes the transition from the DLOR to the VMNR is gradual^29^). Additionally, the VMNR recording might be confounded by the volume conductance of DLOR and/or cortex activity.

**Figure 5:**
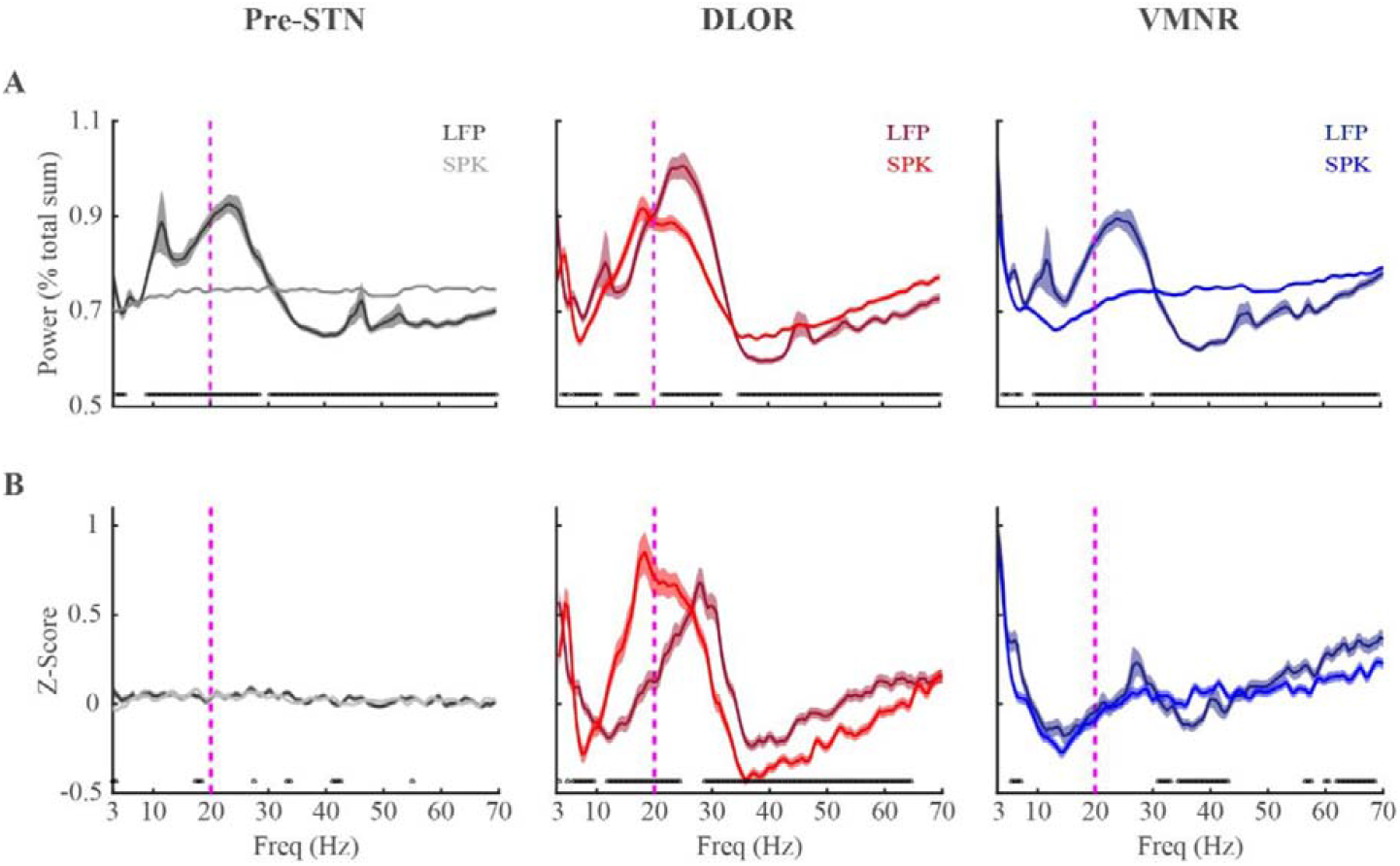
Averaged power spectrum densities show that the peak beta oscillations in spiking (SPK) activity is at a lower frequency than that of LFP in the dorsolateral oscillatory region of the subthalamic nucleus. (A) The PSDs of LFP and SPK are normalized by frequency in three sub-regions. (B) The PSDs of LFP and SPK are normalized by frequency and by distance (Pre-STN activity). The dark and light lines indicate the LFP and SPK respectively in the pre-STN (grey), STN-DLOR (red) and STN-VMNR (blue). Their corresponding shade lines indicate SEM. The black circles above the X-axes indicate frequencies at which there were significant differences (Wilcoxon rank sum test) between LFP and spiking activity. See also Figure S7.

The frequencies of beta oscillations are different across patients. However, they tend to be stable for the same patients and along a single STN trajectory^1^. To further explore the relationship between the frequencies of whitened LFP and whitened SPK beta oscillations, we calculated their beta center frequencies (*β*CFs, referred as LFP *β*CF and SPK *β*CF) for each trajectory. The raster displays of same trajectory LFP and SPK *β*CFs reveal a robust tendency towards the right-lower half (LFP *β*CF > SPK *β*CF, Fig. 6A). LFP *β*CF is significantly higher than SPK *β*CF, and the fraction of pairs whose LFP *β*CF is larger than their corresponding SPK *β*CF far exceeds the fraction that is smaller (Fig. 6A, middle and right subplots). We also estimated *β*CF in PSDs normalized by frequency and distance (i.e., by the pre-STN activity). The SPK *β*CF is relatively downshifted more, and the percentage of LFP *β*CF larger than SPK *β*CF increases after the z-score normalization (Fig. 6B compared to Fig. 6A).

**Figure 6:**
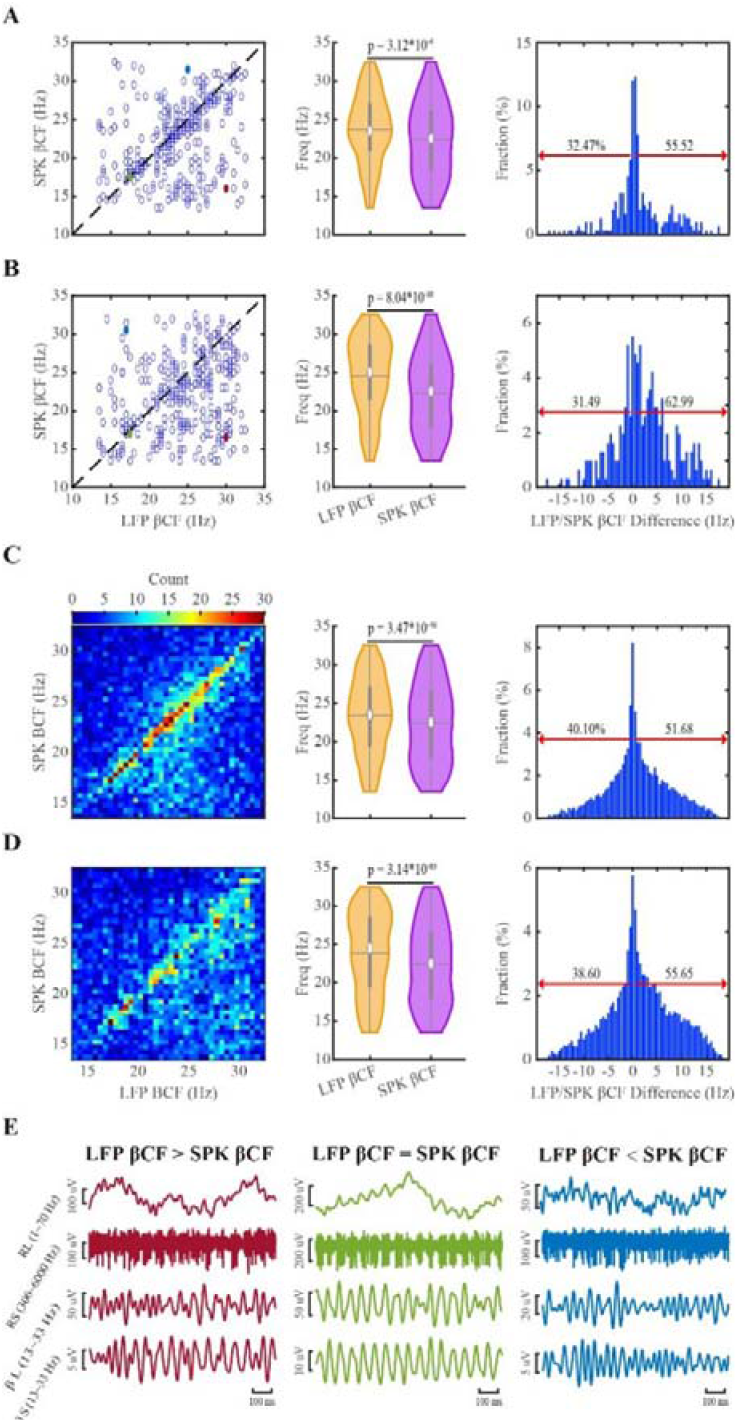
Downshift of the center frequency of beta oscillations of spiking (SPK) activity compared to LFP in the dorsolateral oscillatory region of subthalamic nucleus. (A and B) The unit to get beta center frequencies (βCFs) is trajectory. (C and D) The unit to get βCFs is single recording site. (A and C) The βCFs are obtained from the frequency-normalized power spectra. (B and D) The βCFs are obtained from the frequency- and distance-normalized power spectra. The dark dashed lines on left panel of A and B are the diagonal line (at which x=y). The violins in the middle panel of A, B, C and D demonstrate the distribution of βCFs of LFP and SPK. The significance levels shown in the violin plots were calculated by the Wilcoxon signed rank test. In the right panel, the red arrows indicate the percentage of SPK βCFs that were upshifted (left) and downshifted (right) compared to the corresponding LFP βCFs. (E) shows the raw signals of three examples (LFP βCF is larger than, equal to, or smaller than SPK βCF, from left to right). The examples shown in E are marked in the corresponding colors (red, green, and blue) on the left panel of A and B. RL: raw LFP; RS: raw SPK; βL: β frequency band of LFP; βS: β frequency band of SPK. See also Figures S8, S9 and S10.

Fig. 6C and 6D show *β*CFs from simultaneously recorded LFP and SPK signals of single sites in DLOR of STN calculated from the PSDs normalized by frequency (Fig. 6C) and pre-STN activity (Fig. 6D). At the level of the single recording site (n = 9147), SPK *β*CF also tends to shift downward relative to LFP *β*CF. Whitening in the temporal domain yields similar results (Fig. S8).

### Overlapped distribution and coherence of LFP and spiking activity beta oscillations in DLOR of STN

We calculated the regular and whitened magnitude-squared coherence (Fig. S1) of the simultaneously recorded (in the same recording site) LFP and spiking activity to estimate their frequency overlap and synchronicity. The coherence in beta frequency band is higher in STN DLOR than in the other subregions (Fig. S9A and S9B). There is no significant difference between regular and whitened coherences in any of the subregions (Fig. S9B and S9C). Therefore, the distribution of LFP and spiking beta oscillations overlapped in DLOR of STN, and the LFP-SPK signals have the strongest coherence in the HBeta domain.

### The broader and asymmetric distribution of population SPK and LFP beta oscillations reflects broader distribution of narrow-band frequencies oscillation of single sites with symmetrical band width

The broad and asymmetric distribution of LFP and SPK beta oscillations (Fig. 5) may reflect different scenarios. It could be the result of many single site oscillations with similar broad and asymmetric PSD (Fig. S10A), or of broad and asymmetric distribution of single sites with narrow and symmetric PSD (Fig. S10B). The finding of the down-shift between LFP and SPK *β*CFs (Fig. 6) is consistent with both scenarios, but would reflect a different physiological mechanism. We therefore calculated the half-band widths and half-side widths at the half-height of the beta peaks in 9147 DLOR sites (from 308 trajectories) where both LFP and SPK beta oscillations were simultaneously detected.

The population half-band widths of LFP are narrower than that of SPK (Fig. 7A). After alignment to the *β*CF, the population half-band widths of LFP and SPK are similar (Fig. 7A-C). LFP has a statistically smaller half-band width of beta oscillation in single sites than SPK (Fig. 7D). However, compared to the difference between non-aligned population half-band widths of LFP and SPK (2.3Hz), the difference between their aligned population half-band widths (0.05Hz) and between their half-band widths in single site (0.3Hz) is much smaller. For both LFP and SPK. the aligned population *β* oscillations reveal symmetric distribution (Fig. 7C, S11A and S11B), while the non-aligned population *β* oscillations show asymmetric distribution (Fig. 5A, 5B, S11A and S11B). In the single recording sites, both LFP and SPK beta oscillations have symmetric left and right flanks of half-band widths (Fig. S11A and S11B).

**Figure 7:**
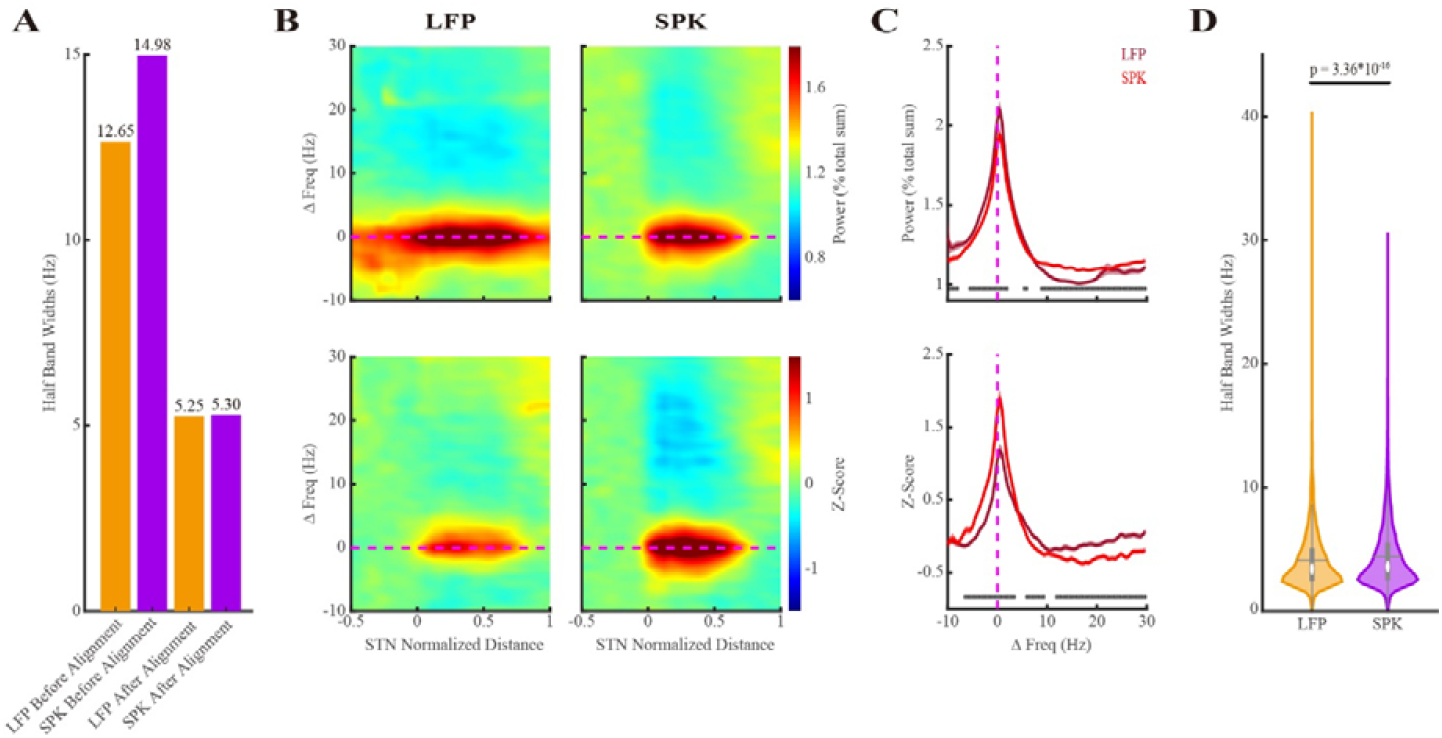
The distributional of beta oscillations of LFP and spiking (SPK) activity in single site is narrower than the population distribution of beta oscillation in the STN DLOR. (A) The population half-band width of LFP and SPK beta oscillations in dorsolateral oscillatory region of subthalamic nucleus. The first and second orange/purple bars indicate the half-band width of LFP/SPK before and after the alignment to the peak beta frequency, respectively. (B) The spectrograms of LFP (left panel) and SPK (right panel) are whitened in the frequency domain and their frequencies are shifted to the peak beta frequency. The color-scale in the first row of B indicates the percentage of total power. The color-scale in the second row of B represents the standard deviation from the mean value of the first 10 depths in pre-STN (z-score). The horizontal magenta dashed line is the reference line of the peak beta frequency (ΔFreq = 0 Hz). (C) The averaged power spectrum of LFP (dark red line) and SPK (light red line) in the dorsolateral oscillatory region (DLOR) the of STN. Their corresponding shade lines indicate SEM. The power spectrum is normalized by frequency (upper subplot) and by frequency and distance (lower subplot). (D) Violin plots of the distribution of half-band widths of LFP and spiking beta oscillations (4.10 ± 2.34 Hz vs 4.40 ± 2.62 Hz (mean ± SD), respectively) in each recording site. The Wilcoxon signed rank test was used for pairwise comparison of half-band widths between LFP and spiking activity. See also Figures S10, S11 and S12.

We also evaluated the 1/4 and 3/4 height band widths and their half-side band widths to confirm that our results are not due to a bias introduced from using only the detection of half-band widths. Similar results were obtained (Fig. S11 and S12). Thus, the different population distribution of LFP and SPK beta oscillations don’t result from the beta frequency distribution of single site (Fig. S10A), but rather reflect different distribution of single sites’ *β*CFs (Fig. S10B). These results therefore indicate that the non-linear transmission of information from LFP to SPK reflects a population downshift of SPK *β*CFs compared to LFP *β*CFs in the STN.

## Discussion

In this study, we highlighted the differences and relationship between LFP and the discharge rate of spiking activity (SPK) in Parkinson’s patients by separating STN neuronal activity into aperiodic and periodic components^23^. We found that the LFP exponent resembled Brown noise (α=2), whereas SPK exponent is close to zero, the exponent of white noise (α=0). In the periodic components, we unexpectedly found that *β*CFs were downshifted in SPK relative to LPF in the motor region (DLOR) of STN. The *β*CFs shift was not caused by a shift in asymmetric distribution of LFP and/or SPK beta oscillations. Rather, our results point to a different distribution of symmetric, narrow oscillations of STN LFP and SPK activities.

### Power-law behavior of subthalamic LFP activity

We found that the LFP displays power-law behavior. The power is inversely and linearly related to the frequency in log-log plots, i.e., there is 1/f^α^ scaling of the power (where α is what we refer as the exponent). We found that LFP exponent in pre-STN is larger than the exponents in STN, but the exponents in DLOR and VMNR don’t differ from one another.

The exponent can be affected by many factors^14^, one of which is the relative contribution of excitation and inhibition (E/I ratio)^24, 30^. However, detailed quantitative anatomy of the relative number and their somatic/dendritic location of STN synaptic input is still missing^31, 32^. Additionally, the E/I balance reflects the physiological efficacy of the synaptic inputs, which is significantly affected by the frequency and pattern of discharge of the GPe^33^, and probably of cortico-STN neurons. Thus, our results showing different exponent values in pre-STN and STN domains cannot be easily framed with the suggested relationships with E/I ratio.

Our results possibly can be explained by the degree of neuronal expenditure. Greater neural expenditure causes flatter slopes (smaller exponent)^34, 35^. In PD, the activation of the basal ganglia is profoundly altered, and STN activity is significantly elevated ^15^. We therefore expect LFP exponent in STN to be smaller than that in Pre-STN. There are other possible explanations for our results as well, and future studies should explore the neuronal/metabolic correlates of the exponent to address this question.

### Power-law behavior of subthalamic SPK activity

For the spiking activity, we filtered the raw signals with the 300-6000Hz bandpass filter. In line with our physiological navigation algorithms^27^, we then rectified the spiking activity by the absolute operator^26^ resulting in a signal indicating the neuronal discharge rates of the multi-unit, or background activity recorded by our electrodes. This is different from most previous studies that have used the spiking activity (e.g., 300-3000 Hz) of well isolated single neurons^36, 37^, however, at the price of masking of low-frequency oscillations^25^.

Using the FOOOF algorithm, we found the exponents of SPK are significantly smaller than those of LFP (Fig. 3). SPK exponents are around zero, which resembles the characteristics of a random process (white noise). This is in line with the Poisson like distribution of spiking activity^12^, the tendency to flat spectrum of cortical and pallidal units^38, 39^, and the demonstration that the PSD of the aggregate of spike trains (with Poisson pattern and refectory period) has a flat spectrum, resembling that of white noise^40^.

Probably, the default, background activity of the STN (as of many structures in the nervous system) is random in order to maximize the information capacity of the system, and to maximize the signal-to-noise ratio of the evoked activity. In any case, the possible mechanism and biological significances of the aperiodic parameters of STN spiking activity require further study.

### The periodic behavior of subthalamic LFP and SPK activity

LFP more likely represents slow sub-threshold currents (primarily post-synaptic potentials) of a large neuronal population from a radius of several millimeters and is considered to be a proxy of the ‘input’ to the local neural network^14^. There are three possible origins of LFP beta oscillations in the STN of PD patients: (1) generated within STN through the network functional connectivity and driven by afferent inputs^3, 6, 16, 17^; (2) generated by the STN neurons themselves (intrinsic properties and subthreshold somatic activity)^18^; (3) generated by the volume conductance of LFP from other locations such as the cortex and other massive subcortical structures^41, 42^. LFP and SPK beta oscillations aren’t always simultaneously present in the same recording electrode^41^. In addition, the magnitude of beta oscillations in LFP in the dorsal STN is larger than that in SPK^3, 41^, which is also shown in Figs. 2, 4 and 5. The data presented here support the notion that the LFP (in the range of 3-70 Hz) in STN mainly results from the afferent inputs and volume conductance. Notably, a major fraction of the volume conductance is from the cortex, which is also a major source of STN afferents (the hyper-direct pathway). Thus, there is a significant overlap of these possible sources of STN LFP activity.

We can consider the spiking activity resulting from action potentials of a neuronal structure as reflecting the ‘output’ of the network (since the fraction of interneurons in the basal ganglia structures is minimal)^26, 43^. PD pathologic mechanism may therefore be better understood by exploring the ‘input-output’ or LFP-SPK relationship in the STN. Previous studies indicate that in the STN of PD: (1) the firing of neurons is phase-locked to LFP beta oscillations^3, 18^; (2) the power of LFP is coherent with that of SPK in the beta frequency band^44^; (3) the beta phase of LFP modulates the amplitude of the LFP high frequency oscillations (HFO, 200-500 Hz). However, most of these studies were carried on small number of patients, and their analysis of periodic phenomena might be confounded by the aperiodic components of the STN activity. Finally, these studies are in line with our finding of sizeable fraction of neurons with similar frequency of LFP and SPK oscillations (trajectories/units close to the diagonal in Fig. 6 and S8), and the LFP-SPK coherence (Fig. S9).

Our study shed light on STN input-output question revealing a downshift of the *β*CFs from LFP (input) to SPK (output) (Figs. 2, 4 and 5). This is correct even for simultaneously recorded LFP and SPK in the same microelectrode, after z-score normalization to remove the volume conducted LFP activity (Figs. 5 and 6). The downshifted *β* CFs between SPK and LFP suggest a non-linear input-output transformation of STN beta oscillations. The STN neurons encode and integrate their inputs (LFP activity) from cerebral cortex, thalamus and GPe, and then decode the outcome as their spiking activity. While LFP may be equally affected by all synaptic inputs, the spiking activity is more affected by excitatory synapses, and by synapses that are close to the soma. This is a possible source of the non-linearity that causes the downshift of the *β* CFs toward the low-beta range in this input/output encoding/decoding STN process. That is, spikes can be dissociated from LFP. which even happened in the cortex^45^. Finally, we expect that the STN spiking activity which drives the central and output structures of the basal ganglia, rather than LFP beta oscillations, probably underlies the motor, and possibly also the non-motor (e.g., sleep) symptoms of PD.

### Conclusions and Limitations

The LFP and spiking activity in the STN of 146 PD patients were separated into periodic and aperiodic components using FOOOF algorithm. We found the exponent of LFP resembled Brown noise and the exponent of the discharge rate (SPK) was similar to white noise. We also found that the *β*CF in DLOR of STN is downshifted in SPK compared to LFP. This downshift wasn’t caused by asymmetrical distribution of beta oscillations in a single STN recording site, and probably reflects the unique input-output relationships of STN neurons. Future studies should test if this shift plays a crucial factor in the development of motor and non-motor impairments in PD patients.

There are several caveats in our study worth noting. Firstly, this is a single center study. Secondly, the results were obtained from Parkinson’s patients. There are no signals from healthy individuals as a control group. Third, it’s difficult to distinguish between power law and log-normal behaviors based on our limited (one-two orders) frequencies (x-scale) tested. Other methods for estimating the exponent, and enabling the detection of other features (e.g. knee) in the log-log plots were not tested here^46^. Finally, the frequency range tested started at 3Hz, and lower frequency (Delta) oscillations were not included. However, the large number of patients and recording sites used in this study support the validity of STN non-linear input-output relationship. This improved understanding of STN pathophysiology and the LFP-SPK beta downshift biomarker will likely pave the way for better adaptive DBS therapy.

## Methods

### Patients

Patients with PD underwent DBS implantation in the STN during the years 2016-2021 at Hadassah Medical Center in Jerusalem, Israel. The patients had to be off medications starting the night before the DBS surgery. Inclusion criteria included clinically established PD, eligibility for DBS procedure, and available intraoperative electrophysiological data in the STN. Additionally, these patients consented to the operative procedure and signed informed consent. This retrospective study was approved by the local Institutional Review Board (IRB) committee (0339-21-HMO).

### Electrophysiological Recordings

Data were acquired with NeuroOmega systems (Alpha Omega Engineering, Ziporit Industrial Zone, Nof HaGalil, Israel). In each hemisphere, 2 microelectrodes (Alpha Omega Engineering) were simultaneously inserted along the planned trajectory targeting the STN in the central and posterior Ben Gun positions, 2mm apart. In rare cases, only one microelectrode was used in the central position due to the anatomy of the patient. All signals were recorded while the patients were awake, at rest, and off medications (overnight washout). The raw signal was sampled at 44 kHz and band-passed from 0.07 to 9,000 Hz using a hardware 2 and 3 pole Butterworth filter, respectively. We began recording at 10mm above target, lowering the electrode between 100-400μm and recording for 4 seconds, after 2 seconds of stabilization, at each site, until we exited the STN. Further details on microelectrode recordings and data acquisition can be found in our previous papers^27^.

### Trajectory selection

308 trajectories were included for analysis out of a total of 492 microelectrode trajectories recorded during the relevant period (January 2016-June 2021). The selection criteria included: (1) the chosen trajectory contained the pre-STN, the dorsal lateral oscillatory region (DLOR) and the ventromedial non-oscillatory region (VMNR); (2) each subregion was longer than 1mm in length. The results reported here were also similar when only the implanted leads were kept in one trajectory per hemisphere (n = 225 trajectories).

### Data Analysis

#### Signal pre-processing

The LFP and spiking signals were obtained by processing the raw data offline (Fig. 1). The LFP was obtained by applying a zero-phase digital 4^th^ order band-pass Butterworth filter with cutoff frequencies 3-200Hz (MATLAB R2020b) to the raw signal. To obtain the spiking signal, we applied a zero-phase digital 4^th^ order band-pass Butterworth filter of 300-6000 Hz. Following this step, we rectified the spiking signal by applying the absolute operator and then subtracted the mean (of the rectified signal).

#### Normalized root mean square (NRMS)

For each recording depth, the RMS of both the LFP and spiking signals were calculated using equation (1)

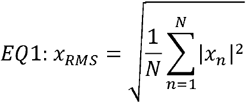

where *x*_*RMS*_ is the RMS value at this site, N is the number of samples in the time domain signal, and *x*_*n*_ is the nth value of the time signal. We normalized the RMS values for each trajectory by dividing each RMS value by the average RMS value of the first 10 sites (presumed to be an unbiased estimation of the baseline activity in the white matter).

#### Power spectral density (PSD)

The PSD was estimated from both LFP and the full wave rectified spiking signal in each recording depth using the pwelch method (MATLAB R2020b), with a Hamming window of 2 seconds (resulting a frequency resolution of 0.5 Hz), 50% overlap and frequency range from 3 to 200 Hz. Any sites with a time signal that was shorter in duration than 1.5 times the window size (i.e., < 3s) was excluded from the analysis. The PSD values of frequencies that are affected by the power-line noise (within 2 Hz of the 50 Hz frequency and its harmonics) were replaced by the mean value of the closest non-affected values. Replacing the values affected by the power-line noise by linear interpolation of the closet values instead resulted in similar results.

The PSD was normalized either by frequency or by frequency and distance. For each recording depth, each PSD value was divided by the total power of the frequency range from 3 to 200 Hz to create a normalized PSD (NPSD). This normalization overcomes the effects of changes in total power (RMS), and will be referred as “normalized by f”. The deviation of NPSD from the mean value of the first 10 depths in pre-STN was calculated, which will be referred as “normalized by f and d” or ‘z-score’.

### Outlier removal

If the value of RMS was more than 3 interquartile ranges above the upper quartile or below the lower quartile, the signal in this recording site was considered to be an outlier and was removed from both the RMS and PSD analyses. The outliers were detected and excluded from the spiking and LFP signals based on their respective RMS.

### FOOOF analysis

The fitting oscillations & one over f (FOOOF) algorithm^23^ was used to separate neural power spectra into aperiodic and periodic components. Aperiodic (offset and exponent) and periodic (center frequency, power, and bandwidth) features were extracted from the LFP and rectified spiking (SPK) signals across the frequency range from 3 to 70 Hz. We translated the FOOOF code from Python to MATLAB language. We added one fitting parameter (peak_width_limits_per) to avoid overfitting in high frequency. Each LFP and SPK PSD was fitted with the following settings: peak_width_limits = [0.8, 12], peak_width_limits_per = [0.02, 0], max_n_peaks = 6, min_peak_height = 0.05, peak_threshold = 2, aperiodic_mode = ‘fixed’.

The FOOOF aperiodic components were used in the whitening procedures detailed below. Finally, we used Spearman’s Rho (correlation coefficient) to calculate the relationship between the aperiodic components (offset and exponent) of LFP and SPK in the Pre-STN, DLOR and VMNR.

### Whitening procedures

In each recording site, the PSD values from 3 to 70 Hz were whitened by multiplying each power by its frequency to the power of alpha as in equation 2:

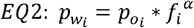

Where 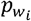 is the whitened power at the i^th^ frequency, 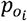is the original power at the i^th^ frequency, *f*_*i*_ is the i^th^ frequency, and α is the aperiodic exponent calculated by applying FOOOF to the PSD data^23^. This whitening method was applied to both LFP and spiking PSD (Fig. 1F). The corresponding whitened PSDs were called “whitened LFP PSD” and “whitened SPK PSD”, respectively. This whitening method will be referred as “whitening in frequency domain (pwelch-FOOOF-whitening)”.

### Time-domain whitening procedure

Classical whitening is done in the frequency domain as detailed in previous section. Here, we also whiten our data in the time domain (https://www.mathworks.com/matlabcentral/fileexchange/65345-spectral-whitening). We applied the time-domain whitening technique to both the LFP and spiking signals (Fig. S1). We first multiplied the time domain signal by an n-point symmetric Hann window (where n is the length of the signal) to diminish spectral leakage. The Fourier transform of this multiplied signal was then obtained with a fast Fourier transform (fft, MATLAB R2020b). The magnitude and phase of each element were extracted from this signal by computing the absolute value and the angle, respectively. The magnitude values, in the frequency range from 3 to 70 Hz, were then whitened by multiplying each magnitude by its frequency to the power of alpha as in equation 3:

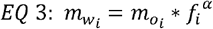

Where 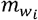 is the whitened magnitude at the i^th^ frequency, 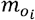 is the original magnitude at the i^th^ frequency, *f*_*i*_ is the i^th^ frequency, and α is the aperiodic exponent calculated from FOOOF on the magnitude data^23^. The modified signal was then transformed back to the time domain using MATLAB’s inverse fast Fourier transform (ifft, MATLAB R2020b) function to obtain the whitened signal (henceforth referred to as the “whitened LFP signal” and the “whitened SPK signal”) (Fig. S1-E1 and S1E2). The whitened PSD was then estimated from the whitened signal in each recording depth using the pwelch method (MATLAB R2020b) as described above. Henceforth, we will refer to this method as whitening in time domain (FFT-FOOOF-whitening-iFFT-pwelch).

### Coherence analysis

The coherence analysis between the LFP and rectified spiking signals of the same microelectrodes was estimated using the magnitude squared coherence function (MATLAB R2020b) with a Hamming window of 2 seconds (for a sampling rate of 44 KHz, resulting a frequency resolution of 0.5 Hz), 50% overlap. We show the frequency range from 3 to 70 Hz (Fig. S1D). We performed the same coherence analysis on the whitened LFP and SPK signals using the same parameters except that we limited the frequency range from 3 to 70 Hz already at stage of the time-domain whitening since the value of aperiodic exponent (alpha) was obtained at this range (Fig. S1F).

### Delimitating STN subregions

The Hidden Markov Model (HMM) algorithm was used to automatically detect the pre-STN, DLOR, VMNR and post-STN regions^27^ from the spiking signal. We used these results to define the regions in both the SPK and LFP analyses (Fig. 1E1 and 1E2).

### Averaging the PSD within safe boundaries

The HMM algorithm enforces sharp transitions between regions. To maximize the reliability of our subregion definition, we chose to exclude the 0.5mm nearest to the border of each region thereby establishing “safe boundaries”. Thus, in the DLOR and VMNR we excluded 0.5 mm nearest to the detected borders of both entry and exit, and for pre-STN we excluded the final 0.5mm preceding the exit (Fig. 1E1 and 1E2). We averaged the PSD within the safe boundaries in each subregion. These averaged PSD from 3 to 70 Hz were used for the FOOOF analysis (Fig. 1F).

### *The simulation of aperiodic and periodic components and the test of their effects on the FOOOF fitting R*^*2*^ *and* mean absolute error *(MAE) values*

We simulated (based on the function (y = αx +b)) 3-70 Hz spectra without Gaussian periodic elements using aperiodic exponent (*α*) ranging from -0.25 to 2.25 (Fig. S2 left subplots). The aperiodic offset was set to equal α, in line with our finding of positive linear correlation between the offset and the exponent (Fig. S3). We also simulated spectra with Gaussian periodic elements using the same aperiodic parameters (Fig. S2, right subplots). In this situation, three periodic Gaussian elements with mean, standard deviation and amplitude values of 18 ± 5 Hz and 1.5 log_10_(power), 25 ± 8 Hz and 3 log_10_(power), and 35 ± 5 Hz and 2 log_10_(power), respectively were added. In addition, a random noise with Gaussian distribution was added into each spectrum to achieve mean absolute error (between spectrums with/without Gaussian noise, range from 0.005 to 0.145). FOOOF analysis was applied to those simulated spectra to obtain their R^2^ and MAE values. R^2^ values were transformed using inverse hyperbolic tangent. We repeated the above process for each permutation and combination of offset, and noise 20 times. R^2^ (after hyperbolic arctangent transformation) and MAE values were averaged and their corresponding standard deviations were calculated. The averaged R^2^ values were transformed back using hyperbolic tangent. Only the average values are shown in Fig. S2.

### Simulation of Brown noise, LFP, spikes and β oscillations

We used the dsp.ColoredNoise function (MATLAB 2021a) to generate a Brown noise signal with a length of 8192 samples (simulating a 2 second signal with a sampling rate of 4096 samples per second). We applied a high-pass 2nd order Butterworth filter with a 0.1Hz cutoff to imitate our patient data which are hardware high-pass filtered at this frequency. We then removed the first and last 2048 samples, leaving only the 4096 middle samples in order to avoid filter edge effects. We simulated the *β* signal by creating a 1 second (4096 samples per second) sine wave at 20Hz with an amplitude of 0.5*SD(x) (where x is the Brownian noise signal). We added the *β* signal to the Brownian noise signal to create the *β* modulated signal. We then rectified the *β* modulated signal (i.e., took the absolute value of the signal) and subtracted the mean of the rectified signals (Fig. S5A-D, left).

For the LFP plus spiking activity simulations (Figs S5F-I, left subplots), we defined a high amplitude beta signal, with amplitude of 1.2*SD(x) in order allow us to define a threshold at which the spikes would ride on the beta peaks, rather than being influenced by low frequency activity due to the Brown noise. We added this high amplitude *β* signal to the Brownian noise signal (x) to generate the high amplitude *β* modulated “membrane potential” signal.

We then defined the spike threshold as the 60th percentile of the high amplitude beta signal, and in the regions where the amplitude of the high amplitude *β* modulated signal exceeded the threshold, we added simulated spikes. The added spikes follow a Poisson distributed probability with a mean of 6 spikes per beta peak. The spike signal was defined as a vector of zeros of the same length as the original signal, with zeros replaced by ones at the time stamps where spikes were generated. The spikes were multiplied by 3*max (high amplitude beta signal) and added to the high amplitude beta modulated brown noise signal. This signal was then band-pass filtered with a 6th order Butterworth filter at 300-2000Hz. Finally, the filtered signal was rectified by taking the absolute operator.

The simulation PSDs were obtained by generating 1000 samples of the time domain signals of each type described above, estimating the spectral density of each using periodogram (MATLAB 2021a) with a Hamming window of 1s and NFFT=4096. We performed a log10 transform on the resultant frequency domain signals and frequencies and averaged the results across the 1000 samples. The PSD results are plotted in figure S5A-I on the right.

### Alignment to the beta center frequency

The averaged PSD within DLOR of each trajectory was used to detect the highest peak beta frequency between 13 and 33 Hz as beta center frequency (*β*CF). The frequency of each site of this trajectory was shifted so the *β*CF is 0 Hz. This alignment enables us to illustrate the relative distribution of power relative to the center frequency. The alignment to the *β*CF was applied to LFP and SPK PSDs, as well as their coherence (Fig. 7, S7 and S9). We used the both averaged trajectory PSD and single site PSD. The use of average trajectory PSD enhances the accuracy of the estimate, and since the frequency of beta oscillations along a single trajectory DLOR is highly stable^1^.

### Calculating band widths of beta oscillation

The normalized PSD (NPSD) was used to find the highest beta peak and its location from 13-33 Hz. Half the highest peak prominence (half-highest-prom) was used as the reference height for width measurement. The half-band width was calculated by finding the distance between the half-highest-prom on the left and right flanks of the beta oscillation (Fig. 7). The powers around the half-highest-prom and their corresponding frequencies were used for linear fitting to get the left and right edges of the half-band width. The distance between the left (or right) edge and the location of highest beta peak was called half-band-half-side width (Fig. S11). It was calculated at three levels. At the level of the single site (n = 9147), we calculated the band widths of beta oscillation separately for each site. At the level of a single trajectory (n = 308), we averaged the NPSD within DLOR of each trajectory and calculated the beta band width from the averaged NPSD. At the population level (n = 1), we used the average of the NPSD within the DLOR of all trajectories to calculate the beta oscillation bandwidth. A similar procedure was done for the 1/4 height and 3/4 height band widths and their half side band widths.

### The simulation of oscillations showing the possible scenarios causing the shift between LFP and SPK in the DLOR of STN

We simulated three broad and asymmetric PSDs with the frequency range from 3 to 70 Hz in single sites: the first one was constructed by three Gaussian elements with mean, standard deviation and amplitude values of 18 ± 5 Hz and 4 log_10_(power) 25 ± 8 Hz and 2 log_10_(power), and 35 ± 5 Hz and 1 log_10_(power), respectively; the second one was constructed by three Gaussian elements with mean, standard deviation and amplitude values of 18 ± 5 Hz and 1 log_10_(power), 25 ± 8 Hz and 4 log_10_(power), and 35 ± 5 Hz and 0.5 log_10_(power), respectively; the third one was constructed by three Gaussian elements with mean, standard deviation and amplitude values of 18 ± 5 Hz and 1.5 log_10_(power), 25 ± 8 Hz and 2 log_10_(power), and 35 ± 5 Hz and 1 log_10_(power), respectively (Fig. S10A, left panel). The three broad and asymmetric PSDs were averaged to generate the population PSD (Fig. S10A, right panel).

Three narrow and symmetric PSDs with the frequency range from 3 to 70 Hz in single sites were simulated with Gaussian elements. Their mean, standard deviation and amplitude values are 18 ± 5 Hz and 1.5 log_10_(power), 25 ± 8 Hz and 2 log_10_(power), and 35 ± 5 Hz and 1 log_10_(power), respectively (Fig. S10B, left panel). The three narrow and symmetric PSDs were averaged to create the population PSD (Fig. S10B, right panel).

## Statistical Analysis

Statistical analyses were performed using MATLAB (R2020b). If not specified, the statistics presented were the mean ± standard deviation (SD) and statistical significance was set at p < 0.05. We used the Bonferroni correction to correct for multiple comparisons. We used the Wilcoxon rank sum test to compare the PSD in each frequency point (Figs. 2, 5, 7, S4, S6 and S7) (two-tailed). The Wilcoxon signed rank test was used for pairwise comparison of beta center frequency and half-band width (Figs. 6, 7, S8, S9, S11 and S12) (two-tailed). The N-way analysis of variance was used to analyze the difference in aperiodic parameters between LFP and SPK (Fig 3 and Table S1, S2 and S3).

## Supporting information

Twelve supplemental figures: Figure S1-S12; 3 supplemental tables: Table S1-S3

## Acknowledgement

This study was supported by the grants of the ISF Breakthrough Research program, Grant NO.: 1738/22, Collaborative research center TRR295, Germany. Project number 424778381, Israel-China bi-national scientific foundation (with Mingsha Zhang, BNU), Grant NO.: 3380/20 and the Silverstein foundation (to Hagai Bergman). The scholarship from China Scholarship Council (to Xiaowei Liu). We thank our patients and the members of the clinical teams at the Hadassah Medical Center, Jerusalem, Israel.

## Contributions

H.B., and J.G., designed the research; Z.I. performed the deep brain stimulation surgery. H.A., J.F.L., and Z.I. acquired the data; X.L., S.G., H.B., and J.G. made the analysis and interpretation of data. X.L. made the first draft of manuscript; All authors read and approved the final manuscript

## Data Availability

The data used in this study is available from the corresponding author upon reasonable request.

## Code Availability

The code used in this study is available from the corresponding author upon reasonable request.

## Notes

### Competing Interest Statement

The authors have declared no competing interest.

## Reference

1. Zaidel, A., Spivak, A., Grieb, B., Bergman, H. & Israel, Z. Subthalamic span of β oscillations predicts deep brain stimulation efficacy for patients with Parkinson’s disease. Brain 133, 2007–2021 (2010).

2. Levy, R., Hutchison, W.D., Lozano, A.M. & Dostrovsky, J.O. High-frequency synchronization of neuronal activity in the subthalamic nucleus of parkinsonian patients with limb tremor. Journal of Neuroscience 20, 7766–7775 (2000).

3. Kühn, A.A., et al. The relationship between local field potential and neuronal discharge in the subthalamic nucleus of patients with Parkinson’s disease. Experimental neurology 194, 212–220 (2005).

4. Darcy, N., et al. Spectral and spatial distribution of subthalamic beta peak activity in Parkinson’s disease patients. Experimental Neurology 356, 114150 (2022).

5. Foffani, G., Bianchi, A.M., Baselli, G. & Priori, A. Movement□related frequency modulation of beta oscillatory activity in the human subthalamic nucleus. The Journal of physiology 568, 699–711 (2005).

6. Brown, P. & Williams, D. Basal ganglia local field potential activity: character and functional significance in the human. Clinical neurophysiology 116, 2510–2519 (2005).

7. van Wijk, B., de Bie, R. & Beudel, M. A systematic review of local field potential physiomarkers in Parkinson’s disease: from clinical correlations to adaptive deep brain stimulation algorithms. Journal of Neurology, 1–16 (2022).

8. Feldmann, L.K., et al. Subthalamic beta band suppression reflects effective neuromodulation in chronic recordings. European Journal of Neurology 28, 2372–2377 (2021).

9. Abosch, A., et al. Long-term recordings of local field potentials from implanted deep brain stimulation electrodes. Neurosurgery 71, 804–814 (2012).

10. Gross, R.E., Krack, P., Rodriguez□Oroz, M.C., Rezai, A.R. & Benabid, A.L.J.M.d.o.j.o.t.M.D.S. Electrophysiological mapping for the implantation of deep brain stimulators for Parkinson’s disease and tremor. 21, S259–S283 (2006).

11. Neumann, W.-J., Köhler, R.M. & Kühn, A.A. A practical guide to invasive neurophysiology in patients with deep brain stimulation. Clinical Neurophysiology (2022).

12. Abeles, M. Local cortical circuits: an electrophysiological study (Springer Science & Business Media, 2012).

13. Asanuma, H. The motor cortex (Raven Press (ID), 1989).

14. Buzsáki, G., Anastassiou, C.A. & Koch, C. The origin of extracellular fields and currents—EEG, ECoG, LFP and spikes. Nature reviews neuroscience 13, 407–420 (2012).

15. Deffains, M., et al. Subthalamic, not striatal, activity correlates with basal ganglia downstream activity in normal and parkinsonian monkeys. Elife 5, e16443 (2016).

16. López-Azcárate, J., et al. Coupling between beta and high-frequency activity in the human subthalamic nucleus may be a pathophysiological mechanism in Parkinson’s disease. Journal of Neuroscience 30, 6667–6677 (2010).

17. van Wijk, B.C., et al. Subthalamic nucleus phase–amplitude coupling correlates with motor impairment in Parkinson’s disease. Clinical Neurophysiology 127, 2010–2019 (2016).

18. Scherer, M., et al. Single-neuron bursts encode pathological oscillations in subcortical nuclei of patients with Parkinson’s disease and essential tremor. Proceedings of the National Academy of Sciences 119, e2205881119 (2022).

19. Kass, R.E., Eden, U.T. & Brown, E.N. Analysis of neural data (Springer, 2014).

20. Cohen, M.X. Analyzing neural time series data: theory and practice (MIT press, 2014).

21. Ermentrout, B. & Terman, D.H. Mathematical foundations of neuroscience (Springer, 2010).

22. Bak, P. How nature works: the science of self-organized criticality (Springer Science & Business Media, 2013).

23. Donoghue, T., et al. Parameterizing neural power spectra into periodic and aperiodic components. Nature neuroscience 23, 1655–1665 (2020).

24. Christoph Wiest, F.T., Alek Pogosyan, Manuel Bange, Muthuraman Muthuraman, Sergiu Groppa, Natasha Hulse, Harutomo Hasegawa, Keyoumars Ashkan, Fahd Baig, Francesca Morgante, Erlick A Pereira, Nicolas Mallet, Peter J Magill, Peter Brown, Andrew Sharott, Huiling Tan. The aperiodic exponent of subthalamic field potentials reflects excitation/inhibition balance in Parkinsonism. Elife 12, e82467 (2023).

25. Yao, B., Salenius, S., Yue, G.H., Brown, R.W. & Liu, J.Z. Effects of surface EMG rectification on power and coherence analyses: an EEG and MEG study. Journal of neuroscience methods 159, 215–223 (2007).

26. Moran, A., Bergman, H., Israel, Z. & Bar-Gad, I. Subthalamic nucleus functional organization revealed by parkinsonian neuronal oscillations and synchrony. Brain 131, 3395–3409 (2008).

27. Valsky, D., Marmor-Levin, O., Deffains, M., Eitan, R. & Blackwell, K.T. Stop! Border Ahead: Automatic Detection of Subthalamic Exit During Deep Brain Stimulation Surgery. Movement Disorders 32, 71 (2017).

28. Zaidel, A., Spivak, A., Shpigelman, L., Bergman, H. & Israel, Z. Delimiting subterritories of the human subthalamic nucleus by means of microelectrode recordings and a Hidden Markov Model. Movement disorders 24, 1785–1793 (2009).

29. Moshel, S., et al. Subthalamic nucleus long-range synchronization—an independent hallmark of human Parkinson’s disease. Frontiers in systems neuroscience 7, 79 (2013).

30. Gao, R., Peterson, E.J. & Voytek, B. Inferring synaptic excitation/inhibition balance from field potentials. Neuroimage 158, 70–78 (2017).

31. Maling, N., Lempka, S.F., Blumenfeld, Z., Bronte-Stewart, H. & McIntyre, C.C. Biophysical basis of subthalamic local field potentials recorded from deep brain stimulation electrodes. Journal of neurophysiology 120, 1932–1944 (2018).

32. Terman, D., Rubin, J.E., Yew, A. & Wilson, C. Activity patterns in a model for the subthalamopallidal network of the basal ganglia. Journal of Neuroscience 22, 2963–2976 (2002).

33. Atherton, J.F., Menard, A., Urbain, N. & Bevan, M.D. Short-term depression of external globus pallidus-subthalamic nucleus synaptic transmission and implications for patterning subthalamic activity. Journal of Neuroscience 33, 7130–7144 (2013).

34. Lendner, J.D., et al. An electrophysiological marker of arousal level in humans. Elife 9 (2020).

35. Arnett, A.B., Peisch, V. & Levin, A.R. The role of aperiodic spectral slope in event-related potentials and cognition among children with and without attention deficit hyperactivity disorder. Journal of Neurophysiology 128, 1546–1554 (2022).

36. Plenz, D., et al. Self-organized criticality in the brain. Frontiers in Physics 9, 639389 (2021).

37. Klaus, A., Yu, S. & Plenz, D. Statistical analyses support power law distributions found in neuronal avalanches. PloS one 6, e19779 (2011).

38. Rivlin-Etzion, M., Ritov, Y.a., Heimer, G., Bergman, H. & Bar-Gad, I. Local shuffling of spike trains boosts the accuracy of spike train spectral analysis. Journal of neurophysiology 95, 3245–3256 (2006).

39. Bair, W., Koch, C., Newsome, W. & Britten, K. Power spectrum analysis of bursting cells in area MT in the behaving monkey. Journal of Neuroscience 14, 2870–2892 (1994).

40. Gao, R. Interpreting the electrophysiological power spectrum. Journal of neurophysiology 115, 628–630 (2016).

41. Marmor, O., et al. Local vs. volume conductance activity of field potentials in the human subthalamic nucleus. Journal of neurophysiology 117, 2140–2151 (2017).

42. Wennberg, R.A. & Lozano, A.M. Intracranial volume conduction of cortical spikes and sleep potentials recorded with deep brain stimulating electrodes. Clinical neurophysiology 114, 1403–1418 (2003).

43. Hardman, C.D., et al. Comparison of the basal ganglia in rats, marmosets, macaques, baboons, and humans: volume and neuronal number for the output, internal relay, and striatal modulating nuclei. Journal of Comparative Neurology 445, 238–255 (2002).

44. Weinberger, M., et al. Beta oscillatory activity in the subthalamic nucleus and its relation to dopaminergic response in Parkinson’s disease. Journal of neurophysiology 96, 3248–3256 (2006).

45. Rule, M.E., Vargas-Irwin, C.E., Donoghue, J.P. & Truccolo, W. Dissociation between sustained single-neuron spiking and transient β-LFP oscillations in primate motor cortex. Journal of Neurophysiology 117, 1524–1543 (2017).

46. Gerster, M., et al. Separating neural oscillations from aperiodic 1/f activity: challenges and recommendations. Neuroinformatics 20, 991–1012 (2022).

