## Supplementary material for "Non-linear input-output relationships in the subthalamic nucleus of Parkinson’s patients": Twelve supplemental figures: Figure S1-S12; 3 supplemental tables: Table S1-S3

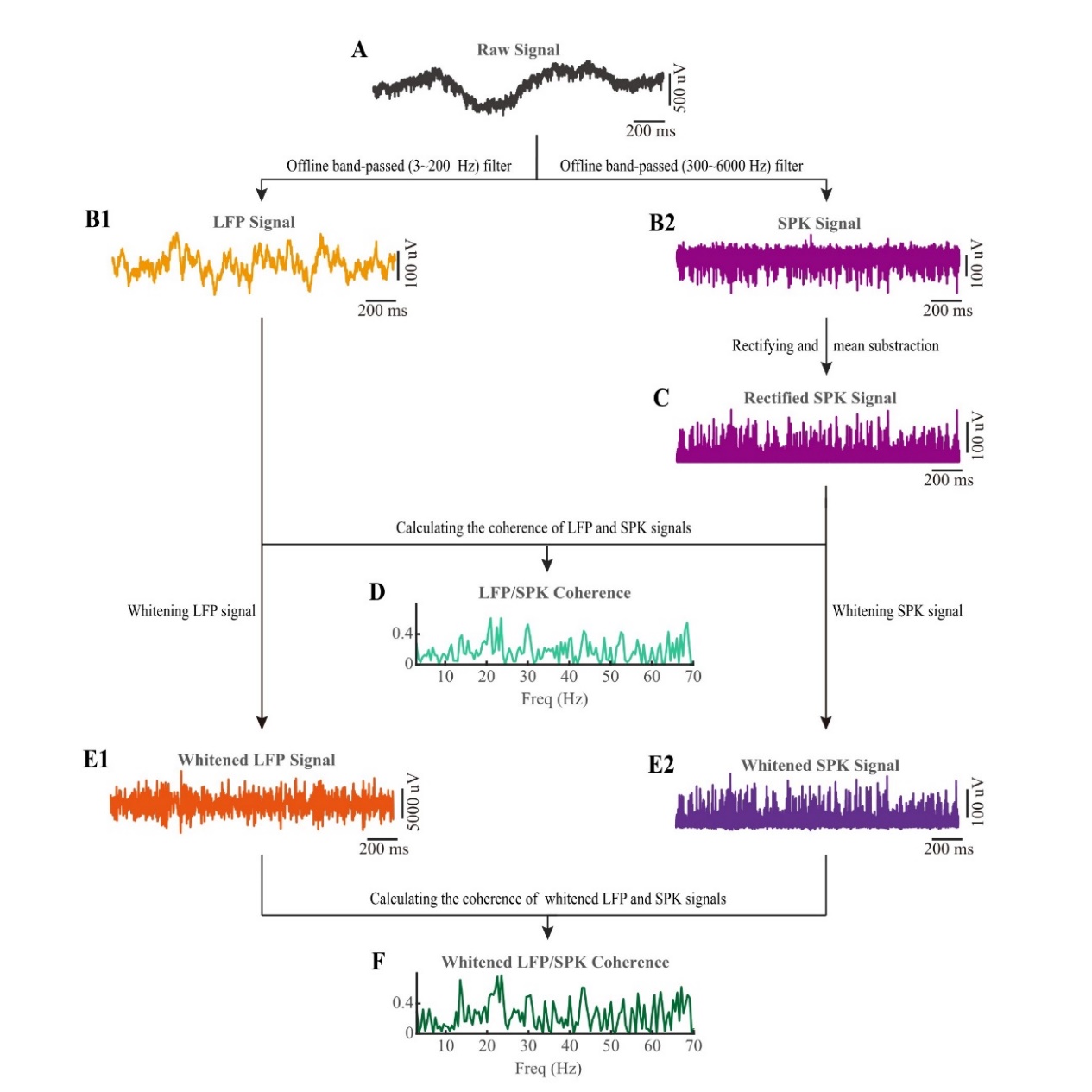

Figure S1: **Calculating the coherence before and after whitening in time domain.** (A) Two seconds example of raw signal band-pass filtered from 0.07 to 9000 Hz. (B1) The signal of LFP from 3 to 200 Hz was obtained by zero-phase filtering of the raw signal. (B2) The signal of spiking activity (discharge rate) was obtained by zero-phase filtering of the raw signal from 300 to 6000 Hz. (C) Applying the absolute operator to the spiking signal and then subtracting the mean created the rectified spiking signal. (D) The coherence of the simultaneously recorded LFP and rectified spiking signals of the same micro-electrode was estimated. (E1 and E2) The whitening technique ($m_{w_{i}}=m_{o_{i}}*{f_{i}}^{\alpha}$; $m_{w_{i}}$ is the whitened magnitude at the i^th ­^frequency, $m_{o_{i}}$ is the original magnitude at the i^th ­^frequency, $f_{i}$ is the i^th ­^frequency, and α is the aperiodic exponent. FFT-FOOOF-whitening-iFFT-pwelch procedure) was applied to the LFP and rectified spiking signals to obtain the whitened LFP signal and the whitened SPK signal. Note the different Y-scale of E1. The whitening process of the LFP (α= $2.20 \pm0.40)$increased the power in high frequency band and amplitude of the LFP signal. (F) The whitened coherence of the simultaneously recorded whitened LFP and SPK signals of the same micro-electrode was calculated. Related to Figure 1.

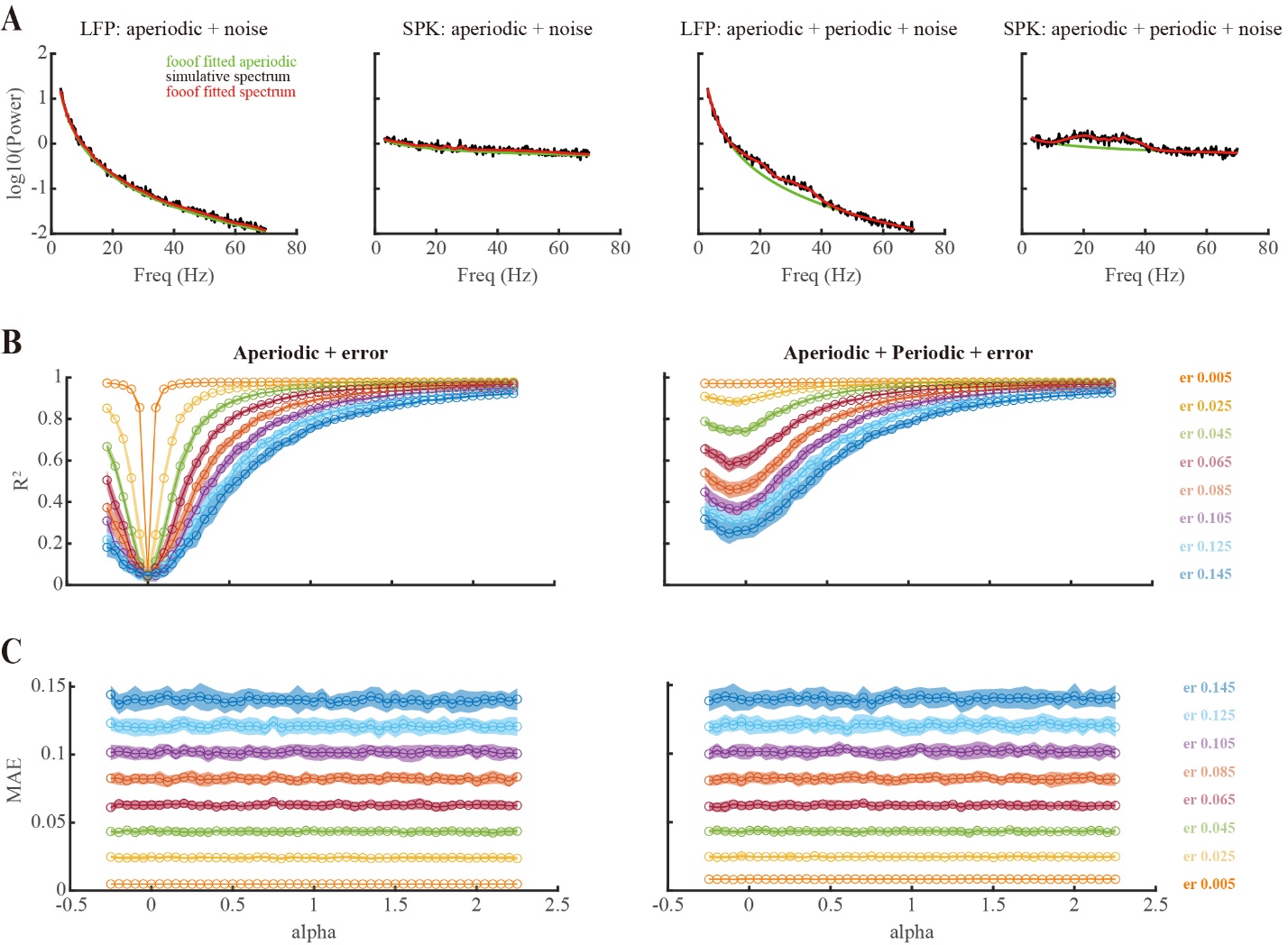

Figure S2: **The absolute value of the aperiodic exponent (α) and periodic oscillations affect fitting R^2^ values but not mean absolute error (MAE).** (A) Examples of simulated aperiodic LFP and SPK spectra (left) and aperiodic plus periodic components spectra (right). (B) In simulated spectra, as the absolute value of $\alpha$ decreases, the value of R^2^ also decreases. The addition of Gaussian periodic elements increases R^2^ values (right) and the addition of Gaussian distributed noise of increasing magnitudes (er, color coded) lowers R^2^ values. (C) α and Gaussian periodic elements don’t change the MAE. As expected, the addition of simulated Gaussian distributed noise of increasing magnitudes increases the MAE. Related to Figures 2 and 3.

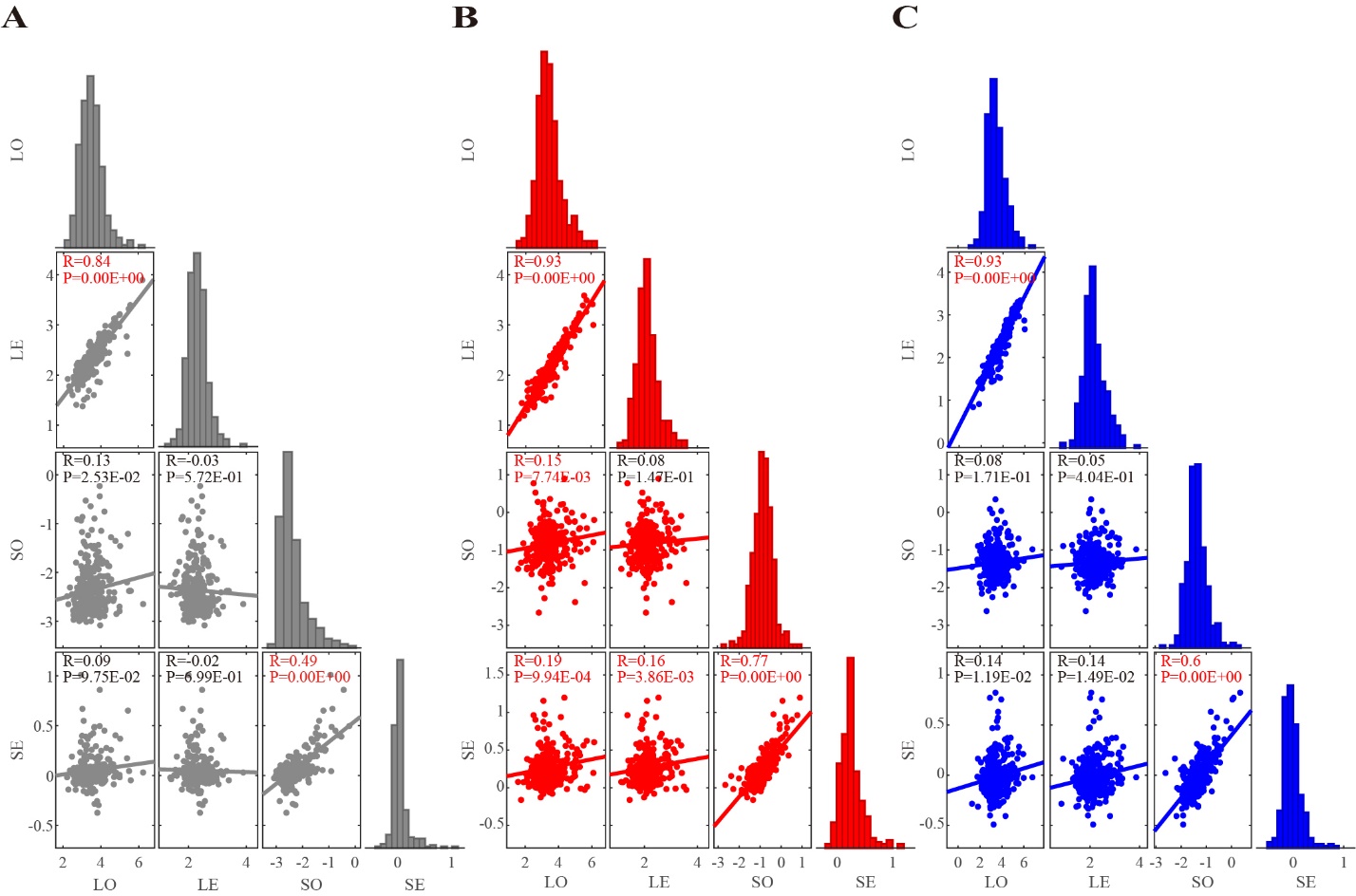

Figure S3: **The correlation between aperiodic parameters of LFP and SPK in three subthalamic subregions.** (A) The correlation between aperiodic parameters of LFP and SPK in the Pre-STN. (B) The correlation between aperiodic parameters of LFP and SPK in the STN-DLOR. (C) The correlation between aperiodic parameters of LFP and SPK in STN-VMNR. LO: LFP offset; LE: LFP exponent; SO: SPK offset; SE: SPK exponent. R indicates the correlation coefficient. P indicates the *P* value for the correlation coefficient to be different from zero. The Pearson’s correlation coefficient was used and the Bonferroni correction was used for significance testing (when p < 0.05/6 = 0.0083, R and P values are typed in red). Plotted along the diagonal of each subplot (A, B, and C) are the distribution histograms of the single parameters. Related to figure 3.

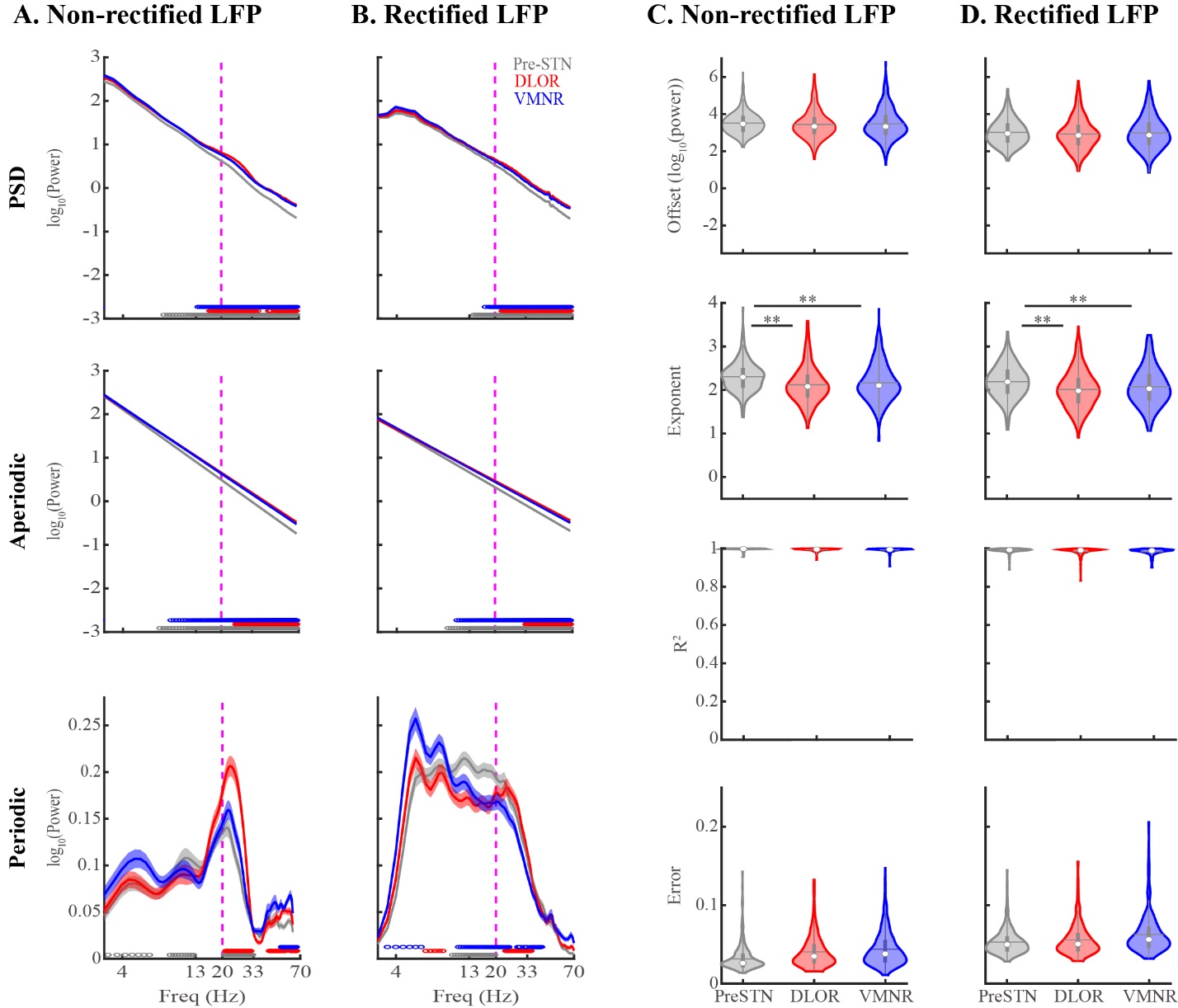

Figure S4: **The aperiodic parameters of non-rectified and rectified LFP are qualitatively similar.** (A) Non-rectified LFP PSD and its aperiodic and periodic components in the pre-STN, STN-DLOR and STN-VMNR sub-regions (grey, red and blue lines, respectively). (B) Rectified LFP PSD and its aperiodic and periodic components in the same sub-regions. Their SEMs are indicated by shade lines with corresponding colors. The circles above the x-axes represent significant differences between the pre-STN and DLOR (grey), the DLOR and VMNR (red), and the VMNR and pre-STN (blue) in corresponding frequency points. We used the Wilcoxon rank sum test and applied the Bonferroni correction. (C) Aperiodic parameters (offset and exponent) and assessment of the goodness of fit (R^2^ and error) of the FOOOF analysis of non-rectified LFP in the three sub-regions. (D) Aperiodic parameters and goodness of fit assessment of the FOOOF analysis of rectified LFP in the three STN sub-regions. Color schemes are the same as in (A, B). The N-way analysis of variance (N-ANOVA) was used to analyze the difference between the aperiodic parameters. The two black asterisks indicate p < 0.01. The detailed multi-comparison of aperiodic parameters (offset and exponent) between non-rectified LFP and rectified LFP is shown in Tables S2 and S3. Related to Figures 2 and 3.

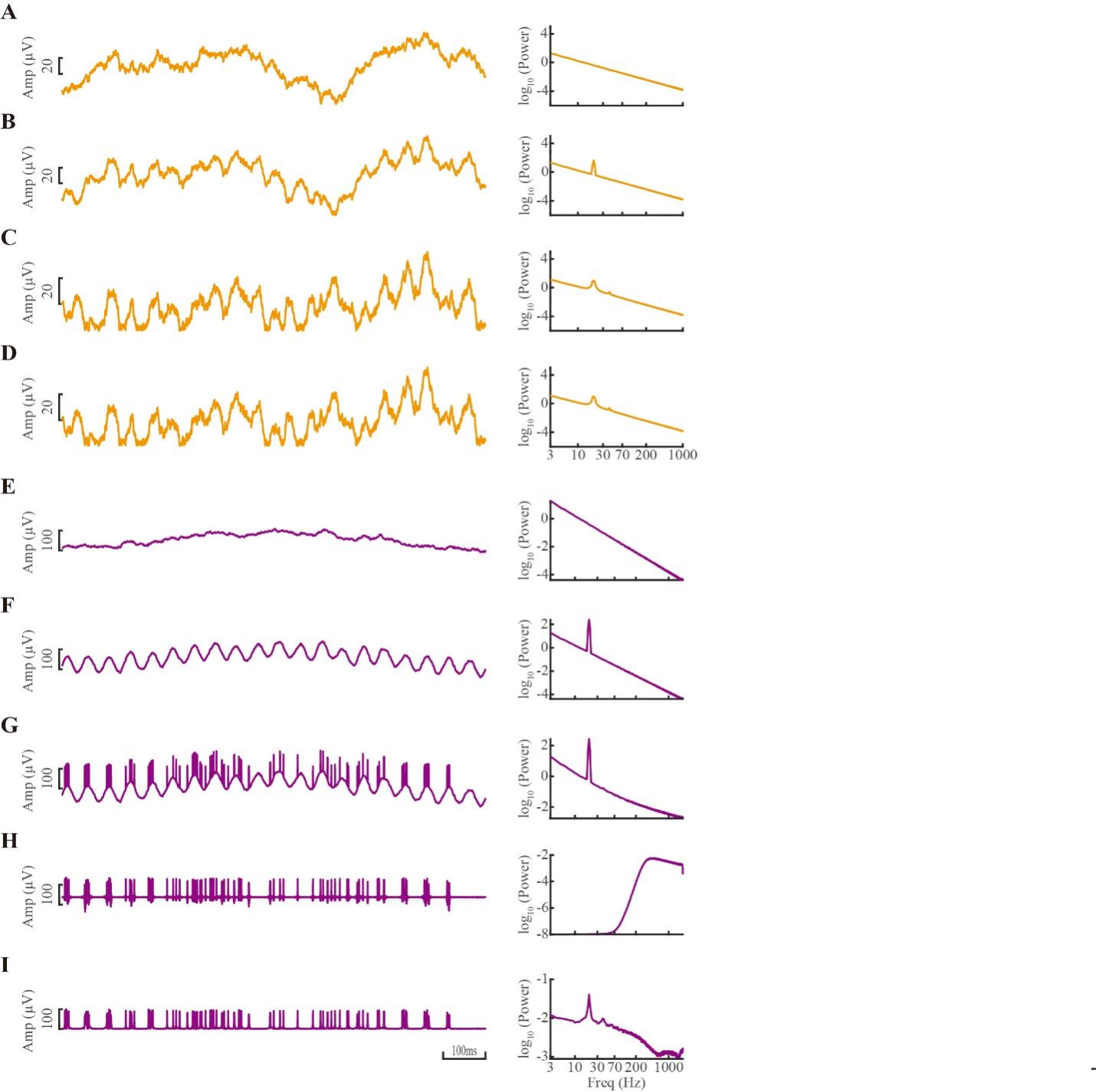

Figure S5: **Simulation of brown noise with beta amplitude modulation of LFP and spiking signals in time (left) and frequency (right) domains.** (A) Brown noise. Left – example of one second trace of the analog time domain signal. Right – power spectral density of 1 second simulation. (B) Brown noise with beta (20 Hz) amplitude modulation. (C) The same as B, but rectified (taking the absolute value of the signal). (D) The same as C, but subtracting its mean value. (E) Brown noise (note the lower resolution of the time-domain Y axis and frequency domain X (Freq) axis compared to A-D). (F) Brown noise and beta (20 Hz) amplitude modulation. (G) Beta modulated Brown noise and Poisson spiking following a threshold crossing. (H) The same as G, but filtered at 300-2000Hz with a zero-phase 6th order Butterworth filter. (I) the same as H, but rectified (i.e., taking the absolute value of the signal and mean subtracted). Related to Figures 2 and 3.

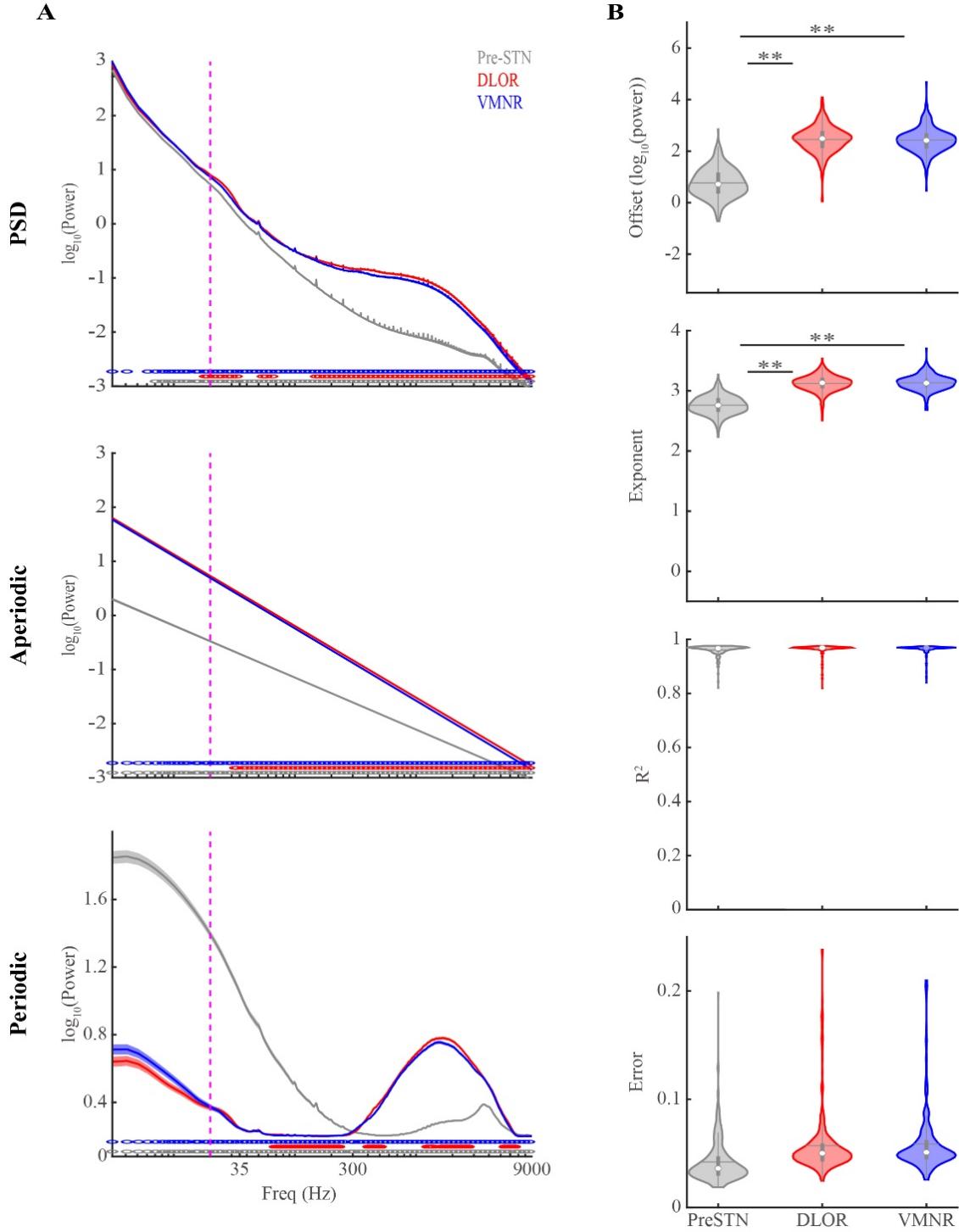

Figure S6: **FOOOF analysis of broad-band (3-9000Hz) raw signal masks low frequency (3-40Hz) periodic oscillations of LFP and spiking activity.** (A) Population average PSD and aperiodic and periodic components (from top to bottom) of raw broad-band signal in three STN sub-regions (Pre-STN shown in grey, DLOR in red, and VMNR in blue). Their SEMs are indicated by shade lines with corresponding colors. The circles above the X-axes represent the significant difference between the pre-STN and DLOR (grey), between the DLOR and VMNR (red), and between the VMNR and pre-STN (blue) in corresponding frequencies. (B) Offset, exponent, R^2^ and error of broad-band raw signal in the three STN sub-regions. The grey/red/blue violins indicate pre-STN, STN-DLOR and STN-VMNR, respectively. The two black asterisks from top to bottom indicate p = 4.28*10^-96^, 2.89*10^-94^, 4.35*10^-90^ and 1.18*10^-86^ (calculated using the Wilcoxon rank sum test and Bonferroni correction). Related to Figures 2 and 3.

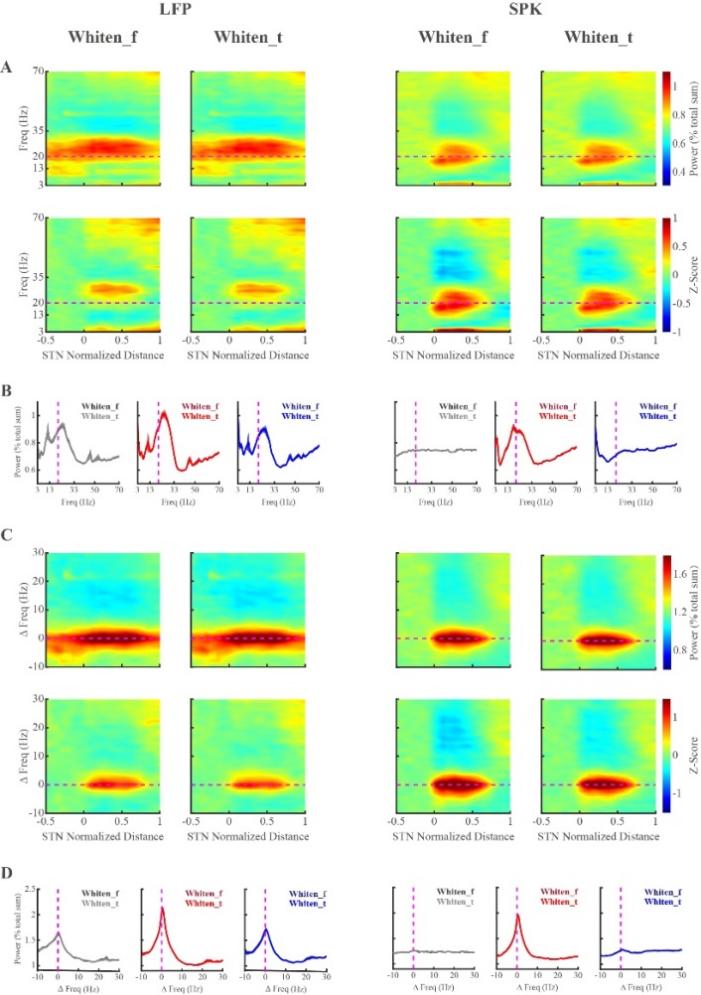

Figure S7: **There is no significant difference in average spectrograms and power spectrum densities obtained with frequency-domain and time-domain whitening method.** The spectrograms and power spectrum densities of LFP and SPK are on the left and right panels, respectively. (A and C) Spectrograms in columns 1 and 3 are whitened in frequency domain and in columns 2 and 4 are whitened in time domain. The frequencies (y axis) in C are aligned to the peak beta frequency. The horizontal magenta dashed lines in A and C are the reference lines of 20 Hz and the aligned peak beta frequency (ΔFreq = 0 Hz), respectively. In the upper row of A and C, the spectrograms are normalized by frequency and the color-scale indicates the percentage of total power. In the lower row of A and C, the spectrograms are normalized by frequency and distance and the color-scale represents the standard deviation from the mean value of the first 10 depths in pre-STN (z-score). (B and D) The LFP power spectrum densities normalized by frequency (i.e. total power in the tested frequency range) are compared between the two whitening methods in three STN sub-regions (the left panel in B and D). The SPK power spectrum densities normalized by frequency are compared between the two whitening methods in three STN sub-regions (the right panel in B and D). The averaged power spectrum densities in B and D are before and after alignment to the peak beta frequency, respectively. The grey/red/blue lines indicate the average power spectrums in pre-STN/DLOR /VMNR whitened in frequency domain (dark) and in the time domain (light), respectively. Their SEMs are indicated by shade lines with corresponding colors. There are no significant differences between the two whitening methods in corresponding frequency points (Wilcoxon rank sum test). Whiten_f: whiten in frequency domain. Whiten_t: whiten in time domain. Related to Figures 4 and 5.

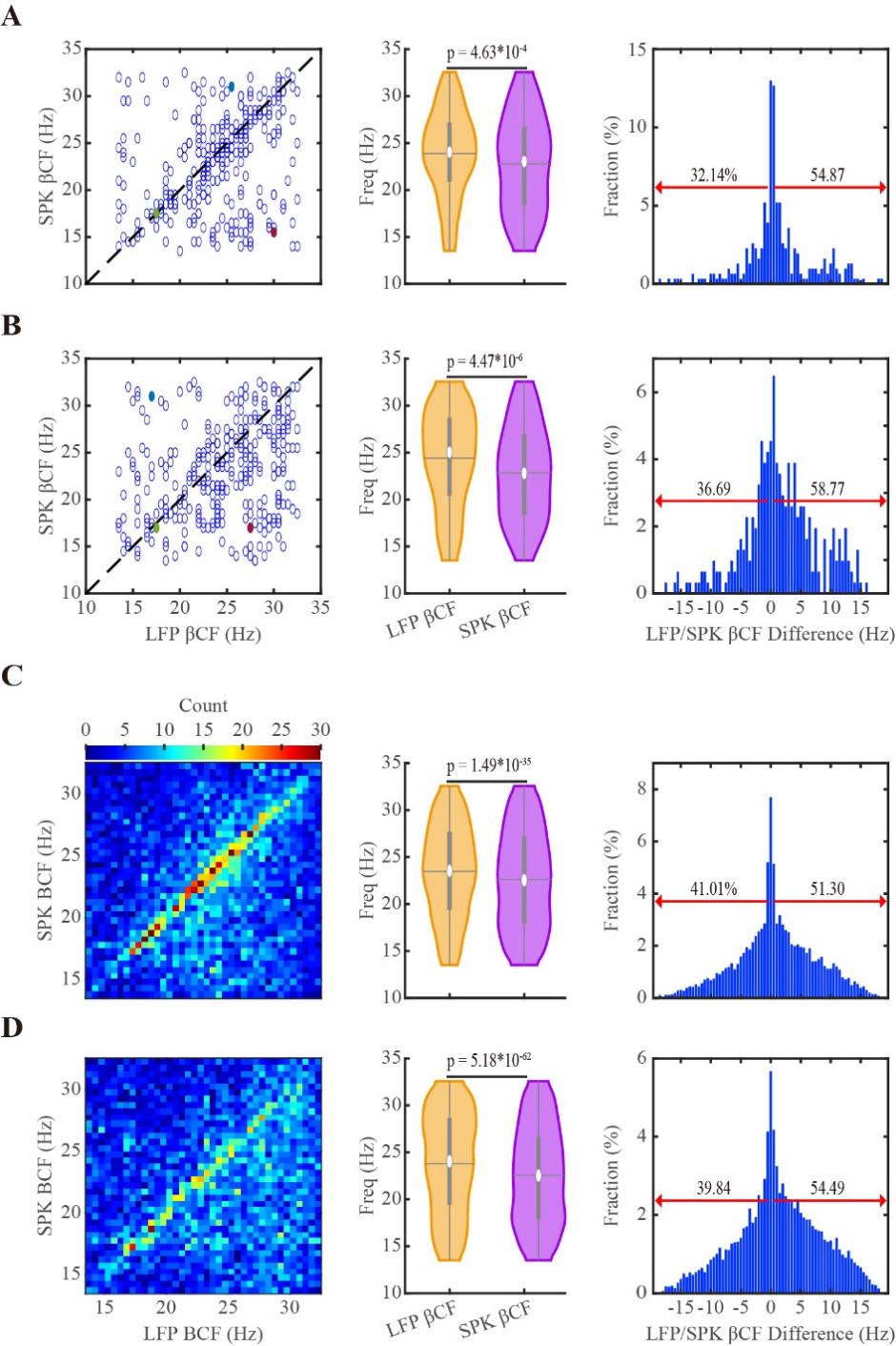

Figure S8: **The downshift of the center frequency of beta oscillations in the time domain whitening method is similar to the downshift detected in the frequency domain whitening method.** (A and B) The unit to get beta center frequencies (βCFs) is trajectory. (C and D) The unit to get βCFs is single recording site. (A and C) The βCFs are obtained from the frequency-normalized power spectra. (B and D) The βCFs are obtained from the frequency- and distance-normalized power spectra. The dark dashed lines in the left column of A and B are the diagonal lines where x=y. In the middle panel, the violins demonstrate the distribution of βCFs of LFP and SPK. The significance levels indicating the difference between the LFP and SPK βCF distributions (shown in the violin plots) were calculated by the Wilcoxon signed rank test. In the right column, the red arrows indicate the percentage of SPK βCFs that were upshifted (left) and downshifted (right) compared to the corresponding LFP βCFs. The filled circles in red, green, and blue in the left column of A and B are the same examples marked in Figure 6. Related to Figure 6.

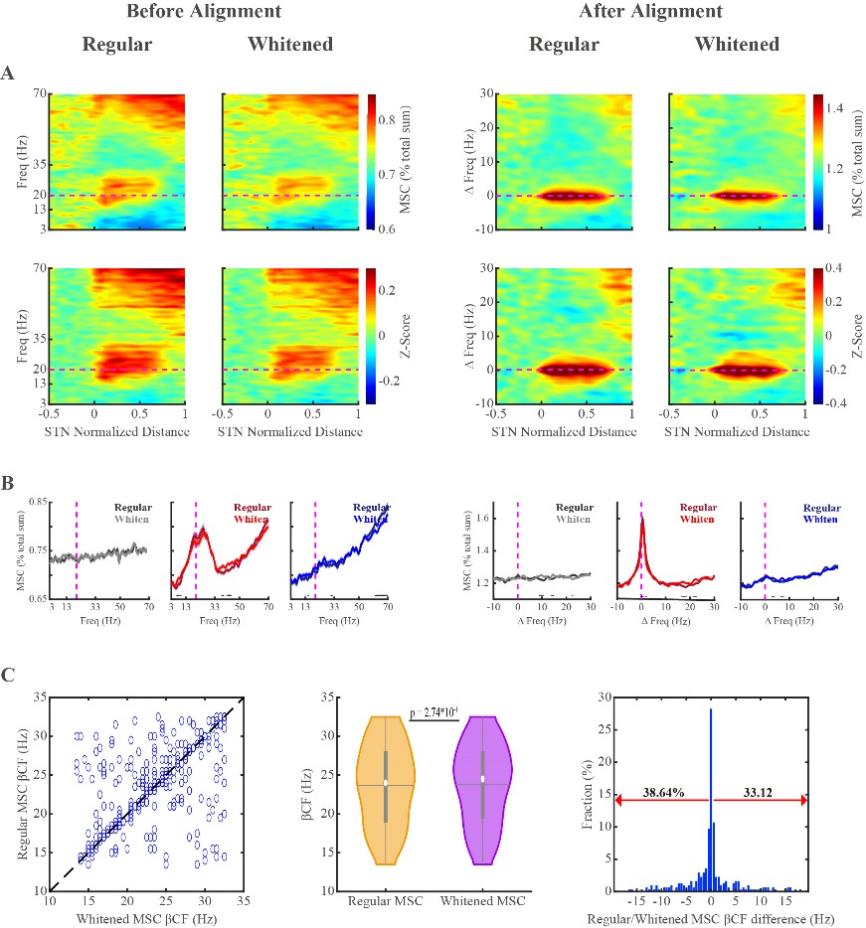

Figure S9: **The regular and whitened coherograms reveal narrow overlap of the distribution of LFP and SPK beta oscillations in the dorsolateral oscillatory region of the subthalamic nucleus.** (A) The coherograms without and with frequency alignment are on the left and right panels of A, respectively. The regular and whitened coherograms before frequency alignment to the peak beta frequency (left panel) are normalized by frequency (top), and by frequency and distance (bottom). The coherograms aligned to the peak beta frequency (right panel) are normalized by frequency, and by frequency and distance (first and second rows), respectively. The horizontal magenta dashed line on the left panel of A is the reference line of 20 Hz. The horizontal magenta dashed line on the right panel of A is the reference line of peak beta frequency (ΔFreq = 0 Hz). The color-scale in the first row of A indicates the percentage of total magnitude-square coherence (MSC). The color-scale in the second row of A represents the standard deviation from the mean value of the first 10 depths in pre-STN (z-score). (B) The comparison of regular and whitened coherence functions in the three STN sub-regions. The dark and light lines indicate the averaged regular and whitened coherence in the pre-STN (grey), DLOR (red) and VMNR (blue), respectively. Their SEMs are indicated by shade lines with corresponding colors. The black circles above the X-axis represent significant difference (Wilcoxon rank sum test) between regular and whitened coherence in corresponding frequencies. (C) The beta center frequencies (βCFs) are obtained from regular and whitened coherence normalized by frequency. The dark dashed line in the left sub-plot is the diagonal line. The Wilcoxon signed rank test was used to calculate the statistical differences between the βCFs of regular and whitened coherence (middle subplot). The red arrows in the right subplot indicate the percentage of βCFs of regular coherence that are smaller (left) or greater (right) than that of the whitened coherence. Related to Figure 6.

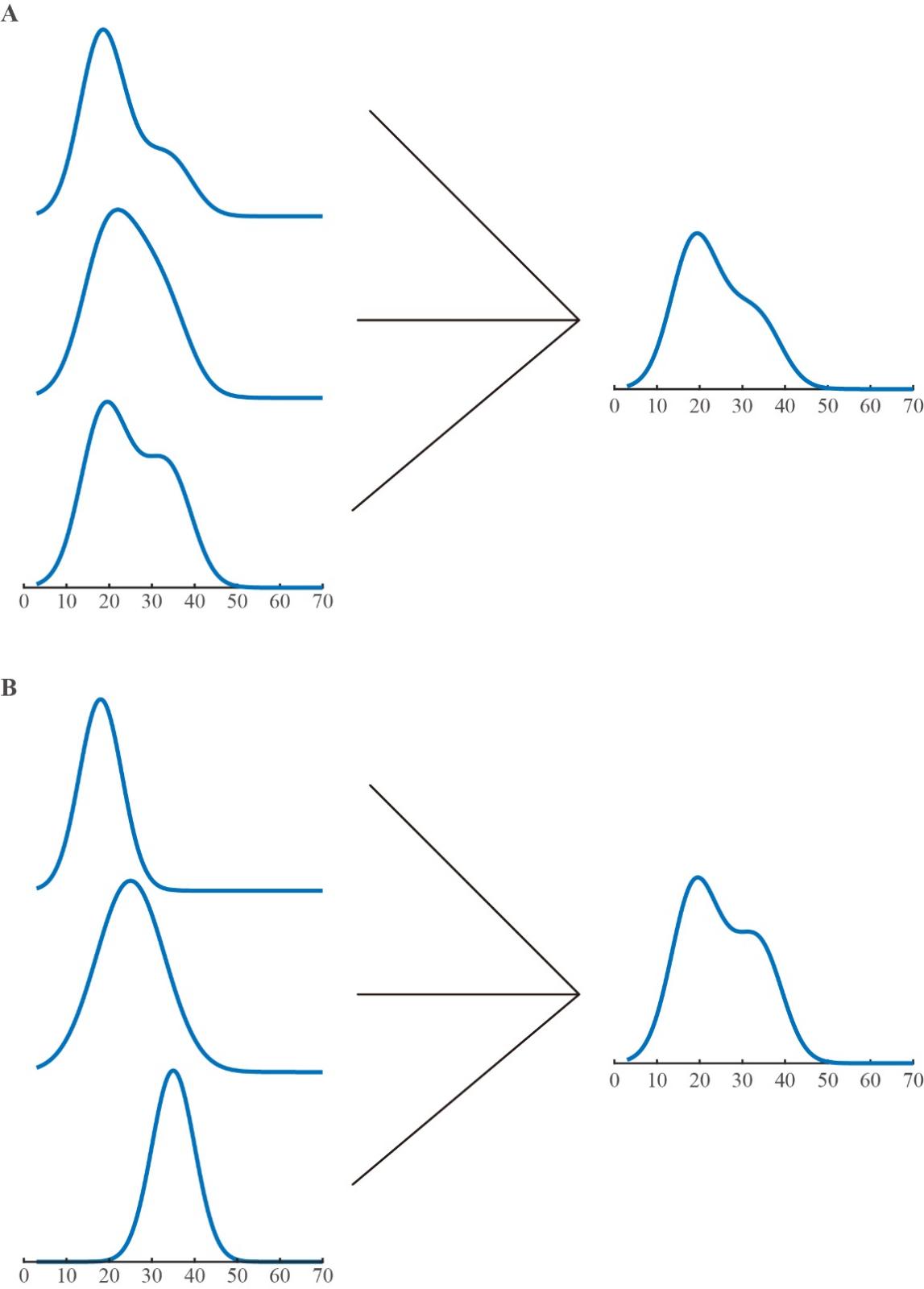

Figure S10: **Two possible scenarios cause the downshift of beta center frequency of SPK relative to LFP in the DLOR of STN.** (A) Three broad and asymmetric power spectral densities in single sites (PSDs) (on the left panel), and similar broad and asymmetric PSD after average of the tree PSDs (on the right panel). (B) Three broad and asymmetric distribution in single sites with narrow and symmetric PSDs (on the left panel), and broad and asymmetric PSD after average of the tree PSDs. Related to Figures 6 and 7.

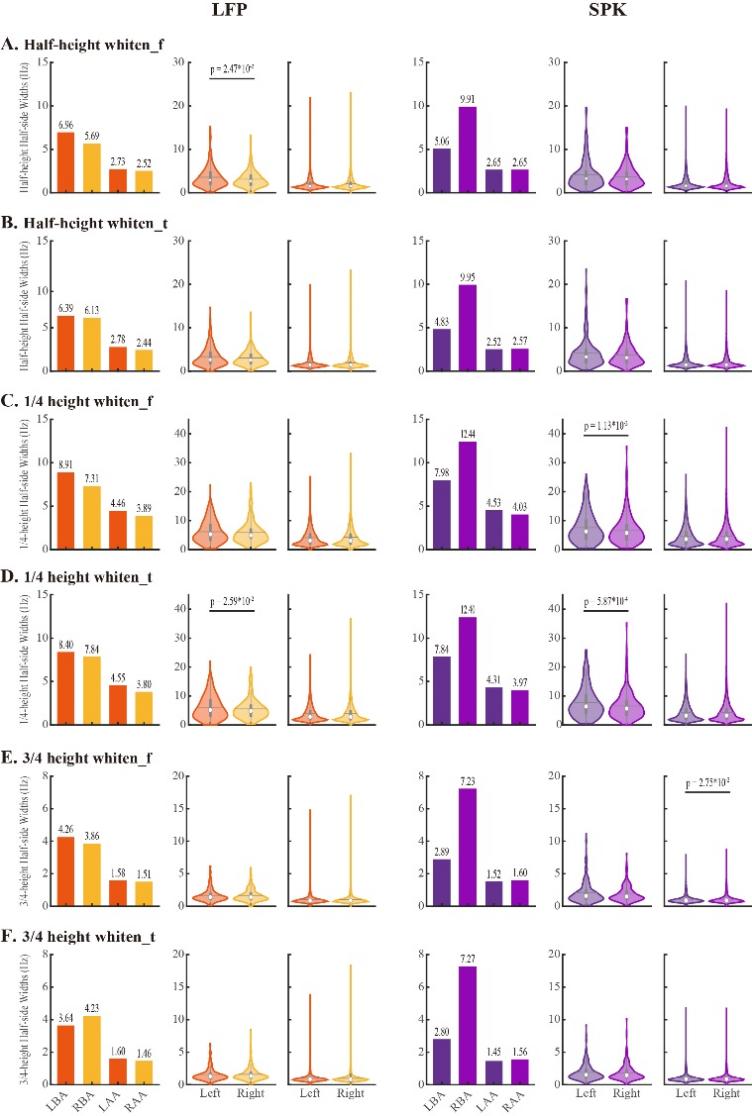

Figure S11: **Symmetrical distribution of left and right half-band widths of beta oscillations of LFP and spiking (SPK) activity in single recording sites but not in the population.** LBA: left before alignment; RBA: right before alignment; LAA: left after alignment; RAA: right after alignment. (A) The left and right half-band widths of LFP and SPK beta oscillations whitened in the frequency domain. (B) The left and right half-band widths of LFP and SPK beta oscillations whitened in the time domain. (C) The left and right 1/4 height half-side band widths of LFP and SPK beta oscillations whitened in the frequency domain. (D) The left and right 1/4 height half-side band widths of LFP and SPK beta oscillations whitened in the time domain. (E) The left and right 3/4 height half-side band widths of the LFP and SPK beta oscillations whitened in the frequency domain. (F) The left and right 3/4 height half-side band widths of the LFP and SPK beta oscillations whitened in the time domain. In the first column of each figure, the first two bars represent the population values without frequency alignment and the last two bars represent the population value with frequency alignment. In the second column, each data point represents the averaged power spectrum in DLOR for each trajectory. In the third column of each graph, each data point represents the power spectrum in a single recording site. The power spectrum used for calculating the beta band widths are normalized by frequency. The left (dark) and right (light) side band widths of LFP and SPK are shown in orange and purple, respectively. We used the Wilcoxon signed rank test to calculate the statistical significance of the differences between right and left half-band widths of beta oscillations. Related to Figure 7.

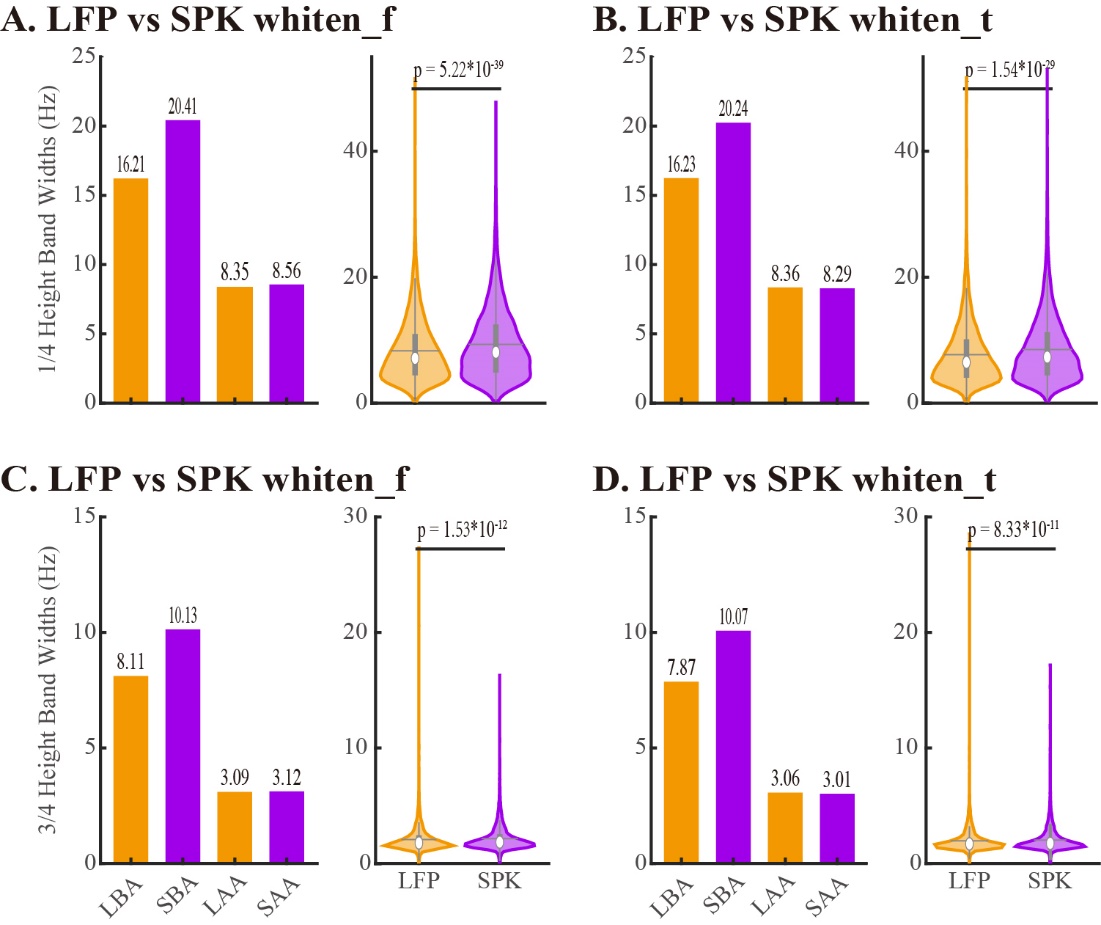

Figure S12: **The comparison of** **both 1/4 and 3/4 height band widths of** $\boldsymbol{\beta}$ **oscillations between SPK and LFP are qualitatively similar to that of half-band width in STN DLOR.**  LBA: LFP before alignment; SBA: SPK before alignment; LAA: LFP after alignment; SAA: SPK after alignment. (A) The two subplots from the left to right demonstrate the comparison of 1/4 height band widths of $\beta$ oscillations between LFP and SPK, which are whitened in frequency domain in population and single site unit, respectively. (B) The two subplots (from the left to right) show the comparison of 1/4 height band widths of $\beta$ oscillations between LFP and SPK which are whitened in time domain in population and single site unit, respectively. (C) The two subplots (from the left to right) reveal the comparison of 3/4 height band widths of $\beta$ oscillations between LFP and SPK which are whitened in frequency domain in population and single site unit, respectively. (D) The two subplots (from the left to right) show the comparison of 3/4 height band widths of $\beta$ oscillations between LFP and SPK which are whitened in time domain in population and single site unit, respectively Related to Figure 7.

| **Table S1: The multi-comparison of aperiodic parameters between LFP and SPK in the three subregions of STN** | | | | | | |
| --- | --- | --- | --- | --- | --- | --- |
| **Parameters** | **Group 1** | **Group 2** | **Lower Limit** | **Mean (Group 1) – mean (Group 2)** | **Upper Limit** | **P value** |
| **LFP vs SPK Offset** | LFP pre-STN | SPK pre-STN | 5.74 | 5.88 | 6.01 | **2.07﹡10^-08^** |
|  | LFP pre-STN | LFP DLOR | -0.06 | 0.07 | 0.21 | 0.65 |
|  | LFP pre-STN | SPK DLOR | 4.20 | 4.34 | 4.48 | **2.07﹡10^-08^** |
|  | LFP pre-STN | LFP VMNR | -0.10 | 0.04 | 0.17 | 0.98 |
|  | LFP pre-STN | SPK VMNR | 4.70 | 4.84 | 4.98 | **2.07﹡10^-08^** |
|  | SPK pre-STN | LFP DLOR | -5.94 | -5.80 | -5.66 | **2.07﹡10^-08^** |
|  | SPK pre-STN | SPK DLOR | -1.67 | -1.54 | -1.40 | **2.07﹡10^-08^** |
|  | SPK pre-STN | LFP VMNR | -5.98 | -5.84 | -5.70 | **2.07﹡10^-08^** |
|  | SPK pre-STN | SPK VMNR | -1.17 | -1.03 | -0.89 | **2.07﹡10^-08^** |
|  | LFP DLOR | SPK DLOR | 4.13 | 4.27 | 4.40 | **2.07﹡10^-08^** |
|  | LFP DLOR | LFP VMNR | -0.18 | -0.04 | 0.10 | 0.97 |
|  | LFP DLOR | SPK VMNR | 4.63 | 4.77 | 4.91 | **2.07﹡10^-08^** |
|  | SPK DLOR | LFP VMNR | -4.44 | -4.30 | -4.17 | **2.07﹡10^-08^** |
|  | SPK DLOR | SPK VMNR | 0.36 | 0.50 | 0.64 | **2.07﹡10^-08^** |
|  | LFP VMNR | SPK VMNR | 4.67 | 4.81 | 4.95 | **2.07﹡10^-08^** |
| **LFP vs SPK Exponent** | LFP pre-STN | SPK pre-STN | 2.18 | 2.25 | 2.32 | **2.07﹡10^-08^** |
|  | LFP pre-STN | LFP DLOR | 0.11 | 0.19 | 0.25 | **2.07﹡10^-08^** |
|  | LFP pre-STN | SPK DLOR | 1.97 | 2.04 | 2.11 | **2.07﹡10^-08^** |
|  | LFP pre-STN | LFP VMNR | 0.07 | 0.14 | 0.21 | **4.56﹡10^-07^** |
|  | LFP pre-STN | SPK VMNR | 2.25 | 2.32 | 2.39 | **2.07﹡10^-08^** |
|  | SPK pre-STN | LFP DLOR | -2.14 | -2.07 | -2.00 | **2.07﹡10^-08^** |
|  | SPK pre-STN | SPK DLOR | -0.29 | -0.21 | -0.14 | **2.07﹡10^-08^** |
|  | SPK pre-STN | LFP VMNR | -2.19 | -2.12 | -2.05 | **2.07﹡10^-08^** |
|  | SPK pre-STN | SPK VMNR | 0.00 | 0.07 | 0.14 | 0.07 |
|  | LFP DLOR | SPK DLOR | 1.79 | 1.86 | 1.93 | **2.07﹡10^-08^** |
|  | LFP DLOR | LFP VMNR | -0.12 | -0.05 | 0.03 | 0.44 |
|  | LFP DLOR | SPK VMNR | 2.07 | 2.14 | 2.21 | **2.07﹡10^-08^** |
|  | SPK DLOR | LFP VMNR | -1.97 | -1.90 | -1.83 | **2.07﹡10^-08^** |
|  | SPK DLOR | SPK VMNR | 0.21 | 0.28 | 0.35 | **2.07﹡10^-08^** |
|  | LFP VMNR | SPK VMNR | 2.11 | 2.18 | 2.25 | **2.07﹡10^-08^** |
| **Mean (Group 1) – mean (Group 2)**: the mean of Group 1 minus the mean of Group 2 is estimated. **Lower Limit**: the lower limit of the 95% confidence interval for the true difference of the means. **Upper Limit**: the upper limit of the 95% confidence interval for the true difference of the means. DLOR：dorsal lateral oscillatory region. VMNR: ventral medial non-oscillatory region. STN: subthalamic nucleus. Related to figure 3. | | | | | | |

| **Table S2: The statistical analysis (N-way ANOVA) between non-rectified LFP and rectified LFP** | | | | | | |
| --- | --- | --- | --- | --- | --- | --- |
| **Non-rectified LFP vs Rectified LFP Offset** | **Source** | **SS** | **DF** | **MS** | **F** | **Prob > F** |
|  | Signal type | 250.00 | 1 | 250.00 | 439.33 | **1.17﹡10^-87^** |
|  | Subregions | 1.96 | 2 | 0.98 | 1.72 | 0.18 |
|  | Signal type*Subregions | 0.02 | 2 | 0.01 | 0.02 | 0.98 |
|  | Error | 1048.19 | 1842 | 0.57 |  |  |
|  | Total | 1300.17 | 1847 |  |  |  |
| **Non-rectified LFP vs Rectified LFP Exponent** | **Source** | **SS** | **DF** | **MS** | **F** | **Prob>F** |
|  | Signal type | 82.36 | 1 | 82.36 | 503.34 | **9.40﹡10^-99^** |
|  | Subregions | 10.47 | 2 | 5.23 | 31.99 | **2.21﹡10^-14^** |
|  | Signal type*Subregions | 0.032 | 2 | 0.016 | 0.098 | 0.91 |
|  | Error | 301.38 | 1842 | 0.16 |  |  |
|  | Total | 394.24 | 1847 |  |  |  |
| **Signal type** indicates statistical analysis of offset (exponent) between LFP and spiking activity. **Subregions** indicates the statistical analysis of offset (exponent) between pre-STN, DLOR and VMNR. **Signal type*Subregions** represents the difference of offset (exponent) between the interaction effect of the signal types and sub-regions. SS: sum of squares due to each source. DF: degree of freedom associated with each source. MS: mean squares for each source. F: F-statistic. Prob > F: the p-value, which is the probability that the F-statistic can take a value larger than a computed test-statistic value. Related to figure S4 | | | | | | |

| **Table S3: The multi-comparison of aperiodic parameters between non-rectified (Non-Rtf) LFP and rectified LFP in the three subregions of STN** | | | | | | |
| --- | --- | --- | --- | --- | --- | --- |
| **Parameters** | **Group 1** | **Group 2** | **Lower Limit** | **mean (Group 1) - mean (Group 2)** | **Upper Limit** | **P value** |
| **Non-Rtf LFP vs Rectified LFP Offset** | Non-Rtf LFP pre-STN | Rectified LFP pre-STN | 0.56 | 0.73 | 0.91 | **2.07﹡10^-08^** |
|  | Non-Rtf LFP pre-STN | Non-Rtf LFP DLOR | -0.10 | 0.07 | 0.25 | 0.83 |
|  | Non-Rtf LFP pre-STN | Rectified LFP DLOR | 0.65 | 0.82 | 0.99 | **2.07﹡10^-08^** |
|  | Non-Rtf LFP pre-STN | Non-Rtf LFP VMNR | -0.14 | 0.04 | 0.21 | 0.99 |
|  | Non-Rtf LFP pre-STN | Rectified LFP VMNR | 0.59 | 0.76 | 0.94 | **2.07﹡10^-08^** |
|  | Rectified LFP pre-STN | Non-Rtf LFP DLOR | -0.83 | -0.66 | -0.49 | **2.07﹡10^-08^** |
|  | Rectified LFP pre-STN | Rectified LFP DLOR | -0.09 | 0.08 | 0.27 | 0.73 |
|  | Rectified LFP pre-STN | Non-Rtf LFP VMNR | -0.87 | -0.70 | -0.53 | **2.07﹡10^-08^** |
|  | Rectified LFP pre-STN | Rectified LFP VMNR | -0.14 | 0.03 | 0.20 | 1.00 |
|  | Non-Rtf LFP DLOR | Rectified LFP DLOR | 0.57 | 0.74 | 0.92 | **2.07﹡10^-08^** |
|  | Non-Rtf LFP DLOR | Non-Rtf LFP VMNR | -0.21 | -0.04 | 0.13 | 0.99 |
|  | Non-Rtf LFP DLOR | Rectified LFP VMNR | 0.52 | 0.69 | 0.86 | **2.07﹡10^-08^** |
|  | Rectified LFP DLOR | Non-Rtf LFP VMNR | -0.96 | -0.78 | -0.61 | **2.07﹡10^-08^** |
|  | Rectified LFP DLOR | Rectified LFP VMNR | -0.23 | -0.06 | 0.12 | 0.95 |
|  | Non-Rtf LFP VMNR | Rectified LFP VMNR | 0.56 | 0.73 | 0.90 | **2.07﹡10^-08^** |
| **Non-Rtf LFP vs Rectified LFP Exponent** | Non-Rtf LFP pre-STN | Rectified LFP pre-STN | 0.34 | 0.43 | 0.52 | **2.07﹡10^-08^** |
|  | Non-Rtf LFP pre-STN | Non-Rtf LFP DLOR | 0.09 | 0.189 | 0.28 | **3.42﹡10^-07^** |
|  | Non-Rtf LFP pre-STN | Rectified LFP DLOR | 0.51 | 0.61 | 0.70 | **2.07﹡10^-08^** |
|  | Non-Rtf LFP pre-STN | Non-Rtf LFP VMNR | 0.04 | 0.14 | 0.23 | **3.75﹡10^-04^** |
|  | Non-Rtf LFP pre-STN | Rectified LFP VMNR | 0.46 | 0.55 | 0.64 | **2.07﹡10^-08^** |
|  | Rectified LFP pre-STN | Non-Rtf LFP DLOR | -0.34 | -0.25 | -0.16 | **2.07﹡10^-08^** |
|  | Rectified LFP pre-STN | Rectified LFP DLOR | 0.08 | 0.18 | 0.27 | **1.03﹡10^-06^** |
|  | Rectified LFP pre-STN | Non-Rtf LFP VMNR | -0.39 | -0.29 | -0.20 | **2.07﹡10^-08^** |
|  | Rectified LFP pre-STN | Rectified LFP VMNR | 0.02 | 0.12 | 0.21 | **4.43﹡10^-03^** |
|  | Non-Rtf LFP DLOR | Rectified LFP DLOR | 0.33 | 0.42 | 0.52 | **2.07﹡10^-08^** |
|  | Non-Rtf LFP DLOR | Non-Rtf LFP VMNR | -0.14 | -0.05 | 0.05 | 0.73 |
|  | Non-Rtf LFP DLOR | Rectified LFP VMNR | 0.27 | 0.37 | 0.46 | **2.07﹡10^-08^** |
|  | Rectified LFP DLOR | Non-Rtf LFP VMNR | -0.56 | -0.47 | -0.38 | **2.07﹡10^-08^** |
|  | Rectified LFP DLOR | Rectified LFP VMNR | -0.15 | -0.06 | 0.03 | 0.46 |
|  | Non-Rtf LFP VMNR | Rectified LFP VMNR | 0.32 | 0.41 | 0.50 | **2.07﹡10^-08^** |
| **Mean (Group 1) – mean (Group 2)**: the mean of Group 1 minus the mean of Group 2 is estimated. **Lower Limit**: the lower limit of the 95% confidence interval for the true difference of the means. **Upper Limit**: the upper limit of the 95% confidence interval for the true difference of the means. DLOR：dorsal lateral oscillatory region. VMNR: ventral medial non-oscillatory region. STN: subthalamic nuclear. **Non-Rtf:** non-rectified. Related to figure S4. | | | | | | |
